# *Aequorea victoria’s* secrets

**DOI:** 10.1101/677344

**Authors:** Gerard G. Lambert, Hadrien Depernet, Guillaume Gotthard, Darrin T. Schultz, Isabelle Navizet, Talley Lambert, Daphne S. Bindels, Vincent Levesque, Jennifer N. Moffatt, Anya Salih, Antoine Royant, Nathan C. Shaner

## Abstract

Using mRNA-Seq and *de novo* transcriptome assembly, we identified, cloned and characterized nine previously undiscovered fluorescent protein (FP) homologs from *Aequorea victoria* and a related *Aequorea* species, with most sequences highly divergent from avGFP. Among these FPs are the brightest GFP homolog yet characterized and a reversibly photochromic FP that responds to UV and blue light. Beyond green emitters, *Aequorea* species express purple- and blue-pigmented chromoproteins (CPs) with absorbances ranging from green to far-red, including two that are photoconvertible. X-ray crystallography revealed that *Aequorea* CPs contain a chemically novel chromophore with an unexpected crosslink to the main polypeptide chain. Because of the unique attributes of several of these newly discovered FPs, we expect that *Aequorea* will, once again, give rise to an entirely new generation of useful probes for bioimaging and biosensing.

## Introduction

EGFP and other engineered variants of avGFP [1] have truly transformed biological imaging, allowing researchers to probe living cells in ways that were previously unthinkable [2–4]. The seemingly impossible task of producing a bright avGFP variant with an emission peak beyond yellow-green [5] drove many groups to explore other marine organisms as potential sources of FPs emitting at longer wavelengths. About five years after avGFP was cloned, FPs were discovered in corals [6,7], and since that time, FPs cloned from jellies, corals, and many other marine organisms have been reported (e.g., [8–10], among many others).

Despite this abundance of reported wild-type FPs, most FPs in widespread use as imaging tools are derived from only a handful of these organisms. Numerous avGFP variants with blue, cyan, green, and yellow-green emission remain the workhorses of livecell imaging, and derivatives of red-emitting FPs from the soft coral *Discosoma* sp. [6,11,12] and the sea anemone *Entacmaea quadricolor* [10,13–15] make up the majority of commonly used FPs emitting at longer wavelengths. With the practical limitations of these particular FP scaffolds becoming more apparent as live-cell microscopy grows more complex and demanding, our group has focused on identifying, characterizing, and engineering FPs with low homology to these traditional choices. Our ongoing efforts include cloning new FPs from diverse sources as well as investigating previously un- or undercharacterized FPs. With this approach, we hope to identify new scaffolds with improved and/or novel spectral properties from which to launch new engineering efforts.

While searching for organisms expressing new and unusual FPs at Heron Island, a research station in the southern Great Barrier Reef, we collected a single individual of an unknown *Aequorea* species that we later determined was most similar to *Aequorea australis*. The striking blue coloration of its radial canals was intriguing (see **Fig. 1**), and led us to reconstruct the transcriptome of the animal. We discovered that, in addition to transcripts encoding an FP clearly homologous to *A. victoria* GFP (avGFP), *A*. cf. *australis* also contained transcripts encoding several other, much more divergent avGFP homologs. We optically characterized the recombinant proteins and solved two of their structures using X-ray crystallography. These investigations revealed a green-emitting FP that is the brightest FP of any color discovered to date, with a molecular brightness nearly 5-fold higher than EGFP. We additionally identified two long-wavelength-absorbing chromoproteins (CPs) with a chemically novel chromophore structure and a reversibly photochromic FP (**Figs. 2** and **3**) in *A*. cf. *australis*.

**Figure 1.**
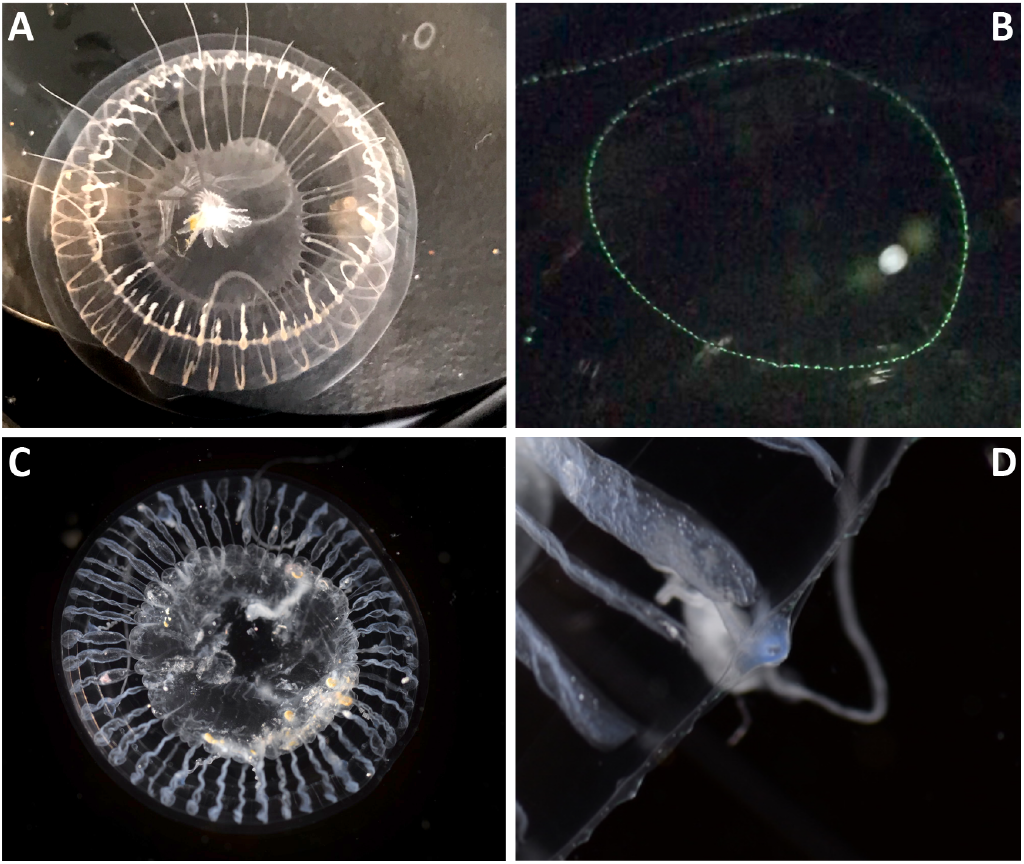
White-light (A) and fluorescence (400 nm LED illumination) (B) photographs of *A. victoria* and white light photographs of *A*. cf *australis* (C, D). The blue coloration of *A*. cf. *australis* is shown in the higher magnification image of one of its tentacle bulbs (D).

**Figure 2.**
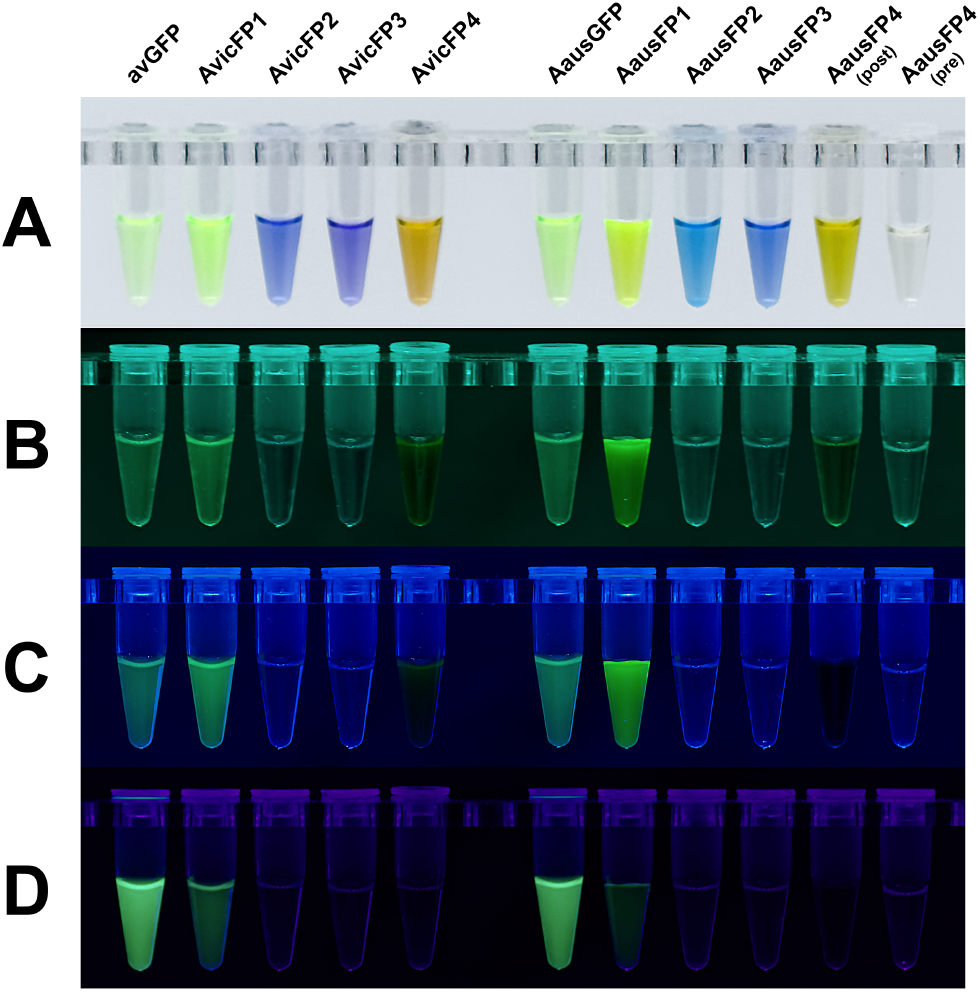
Purified recombinant proteins from *Aequorea* species, shown under (A) white light, (B) 505 nm LED, (C) 480 nm LED, and (D) 400 nm LED illumination. AausFP4 is shown as two tubes containing equivalent protein samples before and after photoconversion by exposure to ~360 nm UV light. All fluorescence photographs were taken without emission filters against a matte black background.

**Figure 3.**
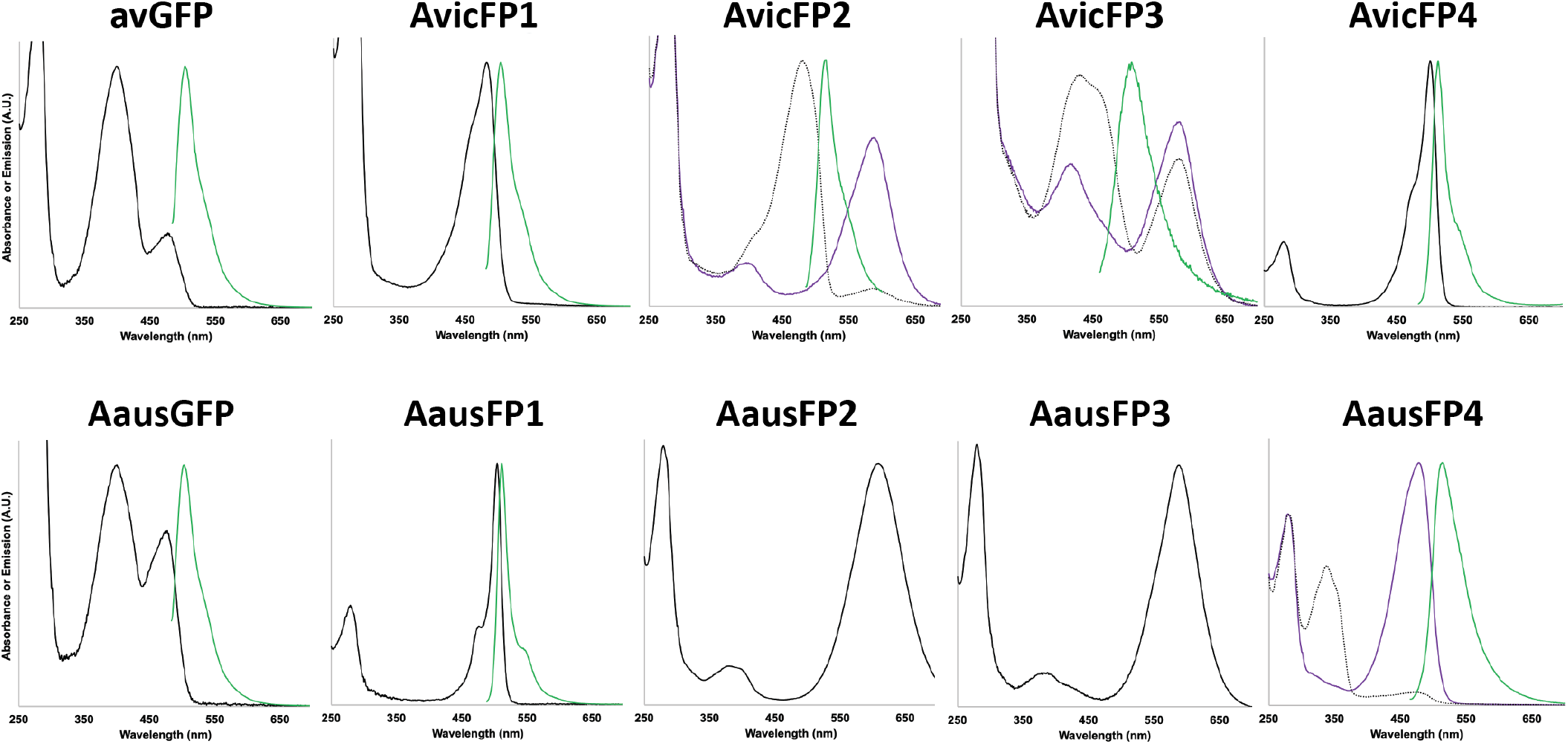
Absorbance and emission spectra (where measurable) for FPs in this study. For photoswitchable and photoconvertible proteins, pre-switching absorbance spectra are shown as doZed lines and post-switching absorbance spectra as solid purple lines. Emission spectra are shown as green lines. The emission spectra for AvicFP2 and AvicFP3 were measured using 460nm excitation prior to photoconversion. The emission spectrum of AausFP4 was measured using 440nm excitation aPer photoswitching to the blue-absorbing state.

Intrigued, we next investigated a sample of *A. victoria* from the Crystal Jelly exhibit at the Birch Aquarium at Scripps to determine whether this species also contained FPs other than avGFP. As suspected, the *A. victoria* individual we sequenced expressed orthologs of the bright green-emitting FP and the unusual CPs that we first identified in *A*. cf. *australis*. Surprisingly, *A. victoria* also expresses a close homolog of avGFP with much-improved properties: a fully anionic chromophore, low pK_a_, and much more efficient folding and maturation at 37°C than wild-type avGFP. Only 2 mutations—one to speed maturation at 37°C and one to monomerize the protein—generated an FP superior to EGFP, the first widely-adopted avGFP variant. *A. victoria*’s two CPs are distinct from those of *A*. cf. *australis*, undergoing a unique mode of photoconversion from a green-emitting (GFP-like) state to a non-fluorescent, red-absorbing state after exposure to low to moderate intensities of blue light (see **Fig. 3**). Evidence so far suggests that this photoconversion chemistry has not been described previously.

## Results and Discussion

### Multiple, Diverse *Aequorea* Green Fluorescent Proteins

As expected, both *Aequorea* species abundantly express close homologs of avGFP. Both **AausGFP** and the **avGFP** identified in this work possess optical and biochemical properties similar to the original avGFP clone [1], characterized by an excitation spectrum with two peaks at 398 and 477 nm and an emission peak at 503 nm. The extinction coefficient and quantum yield of the two proteins are similar, and in our hands match closely with the literature values for avGFP [2].

The first surprise among the newly discovered *A. victoria* GFPs was **AvicFP1**, a transcript with relatively low expression but with high homology to avGFP (80% amino acid identity, see **Fig. 4**). At neutral pH, AvicFP1 has a single absorbance peak at 481 nm, indicating that its chromophore exists in a fully anionic state. Its fluorescence pK_a_ of 4.9, which we attribute largely to the presence of cysteine in the first chromophore position rather than serine in avGFP, is substantially lower than that of EGFP (6.0). The S65T substitution in avGFP is among the most critical early mutations introduced to generate an all-anionic chromophore, though S65C has been reported as well [2,16]. Because of mutations derived from errors in the oligonucleotides used for synthetic gene assembly, we also identified one colony among thousands of the initial AvicFP1 clones that produced a much larger proportion of mature FP in *E. coli* incubated at 37°C. This clone contained a single point mutation leading to the substitution F64L, the second mutation originally identified for avGFP to produce EGFP [2,17], and produces a protein that maintains all of the beneficial properties of the wild-type protein.

**Figure 4.**
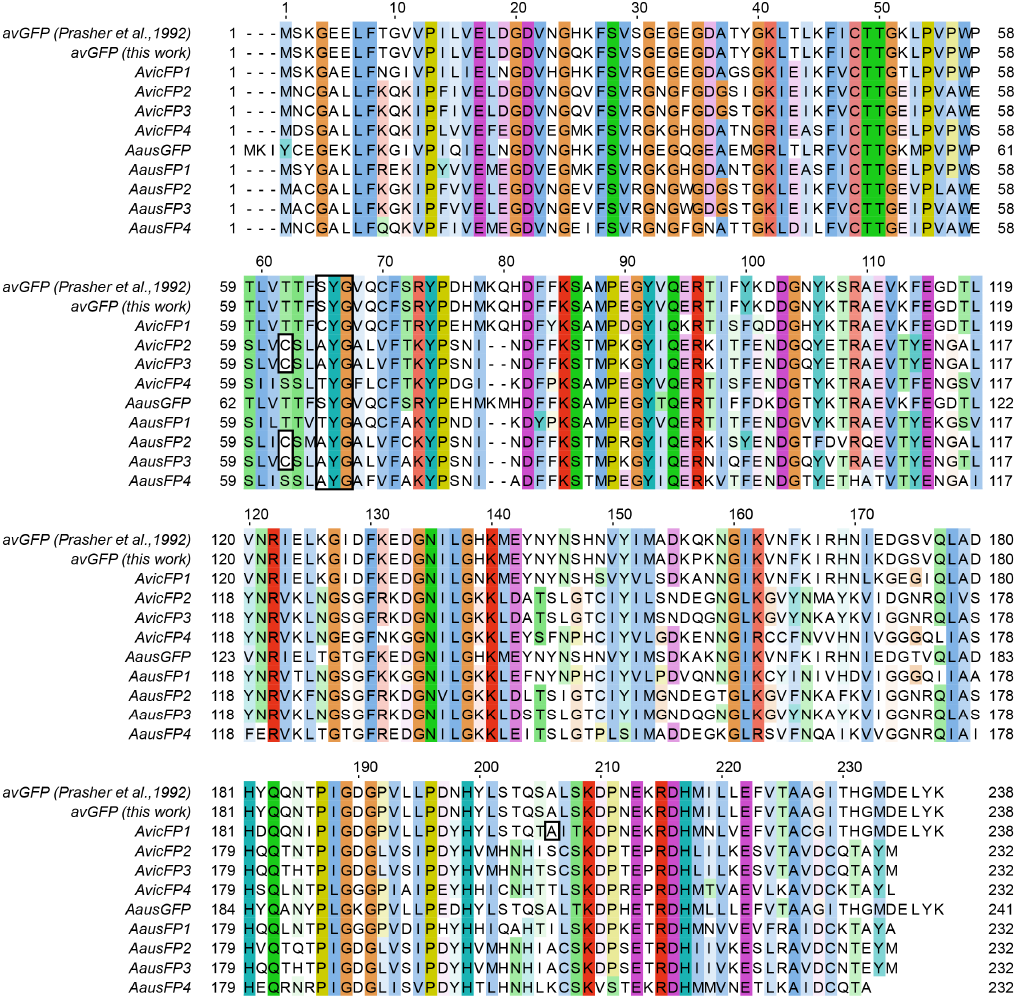
Amino acid sequence alignment of all fluorescent protein homologs identified in *Aequorea* species in this study. Alignment was done using Clustal Omega [25]. Conserved and semi-conserved residues are colored using the CLUSTAL palette. The chromophore tripeptide is outlined by a black box. Additional residues discussed in the text are indicated by additional black boxes. Amino acid residues numbering from wild-type avGFP is given above each line.

A careful examination of the sequence alignment between AvicFP1 and avGFP revealed that essentially all of the side chains that participate in the weak dimer interface of avGFP are conserved in AvicFP1. Based on this observation, we hypothesized that mutations sufficient to monomerize avGFP variants (i.e. A206K [18]) would also produce a monomeric variant of AvicFP1. Using the organized smooth endoplasmic reticulum (OSER) assay to test for oligomeric behavior in cells [19], we found that the mutant AvicFP1-F64L/A206K displays monomeric behavior equivalent to mEGFP, widely considered the “gold standard” of monomeric FPs [19] (OSER data are summarized in **Table S1**). Fusions to LifeAct [20] and histone 2B (H2B) displayed the expected localization (**Fig. 5**), and did not appear to interfere with mitosis or cell growth (qualitative observations only). We therefore named this protein **mAvicFP1**, or “monomeric *A. victoria* fluorescent protein 1.” It is somewhat ironic that avGFP, which required a large number of mutations to fold and mature optimally in mammalian cells, was being expressed in *A. victoria* alongside a much more easily engineered protein. We can only speculate about what biological imaging might look like today if AvicFP1, rather than avGFP, had been the first FP cloned.

**Figure 5.**
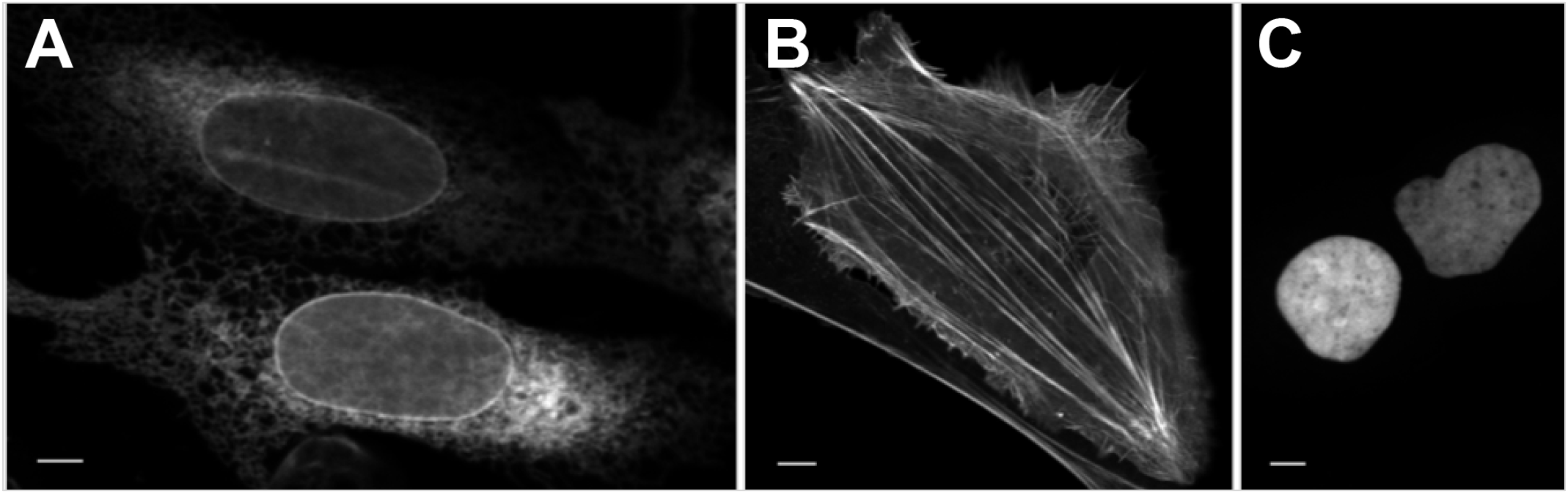
mAvicFP1 fusions to (A) CytERM, (B) LifeAct, and (C) H2B. U2-OS cells display expected localization. Scale bar is 10μm.

We were surprised to discover a second green-emitting FP in *A*. cf. *australis*, **AausFP1**, that shares only 53% amino acid identity with avGFP (see **Fig. 4**). AausFP1 is the brightest FP discovered to date, with a nearly perfect quantum yield (0.97) and a peak extinction coefficient of 170,000 M^−1^cm^−1^, making it nearly fivefold brighter than EGFP on a per-molecule basis. These already extraordinary properties are further bolstered by a low fluorescence pK_a_ (4.4) and unusually narrow excitation and emission peaks (see **Fig. 3**; the emission peak of AausFP1 has a full width at half maximum (FWHM) of 19nm, compared to 32nm for EGFP). Comically, the ortholog of AausFP1 in *A. victoria*, **AvicFP4**, shares some of its unusual properties, such as narrow excitation and emission peaks, efficient folding at 37°C, and a fairly high extinction coefficient, but its low quantum yield (0.10) makes it the dimmest GFP found in *A. victoria*.

AausFP1 was expressed at very low levels relative to other FPs in the *A*. cf. *australis* individual sequenced (see **Supplementary Table S8**), and would be rare or absent in most cDNA expression-cloning libraries. The transcriptomic approach used in this study is the only practical way to identify such unusual, low-abundance FPs, short of costly whole genome sequencing. Despite low expression in its native context, wild-type AausFP1 expresses and folds very efficiently in *E. coli* at 37°C without any modifications. X-ray crystallography revealed a uniquely stabilized chromophore environment in AausFP1 that may be responsible for its unique properties (see **Fig. 6** and **Supplementary Results and Discussion**). Like many wild-type hydrozoan fluorescent proteins, AausFP1 is an obligate dimer. To take advantage of its many advantageous properties, we have optimized tandem-dimer and monomeric variants of AausFP1 for use as fusion tags and biosensor components, and we anticipate that these proteins will be highly useful tools for biological imaging (to be reported separately).

**Figure 6.**
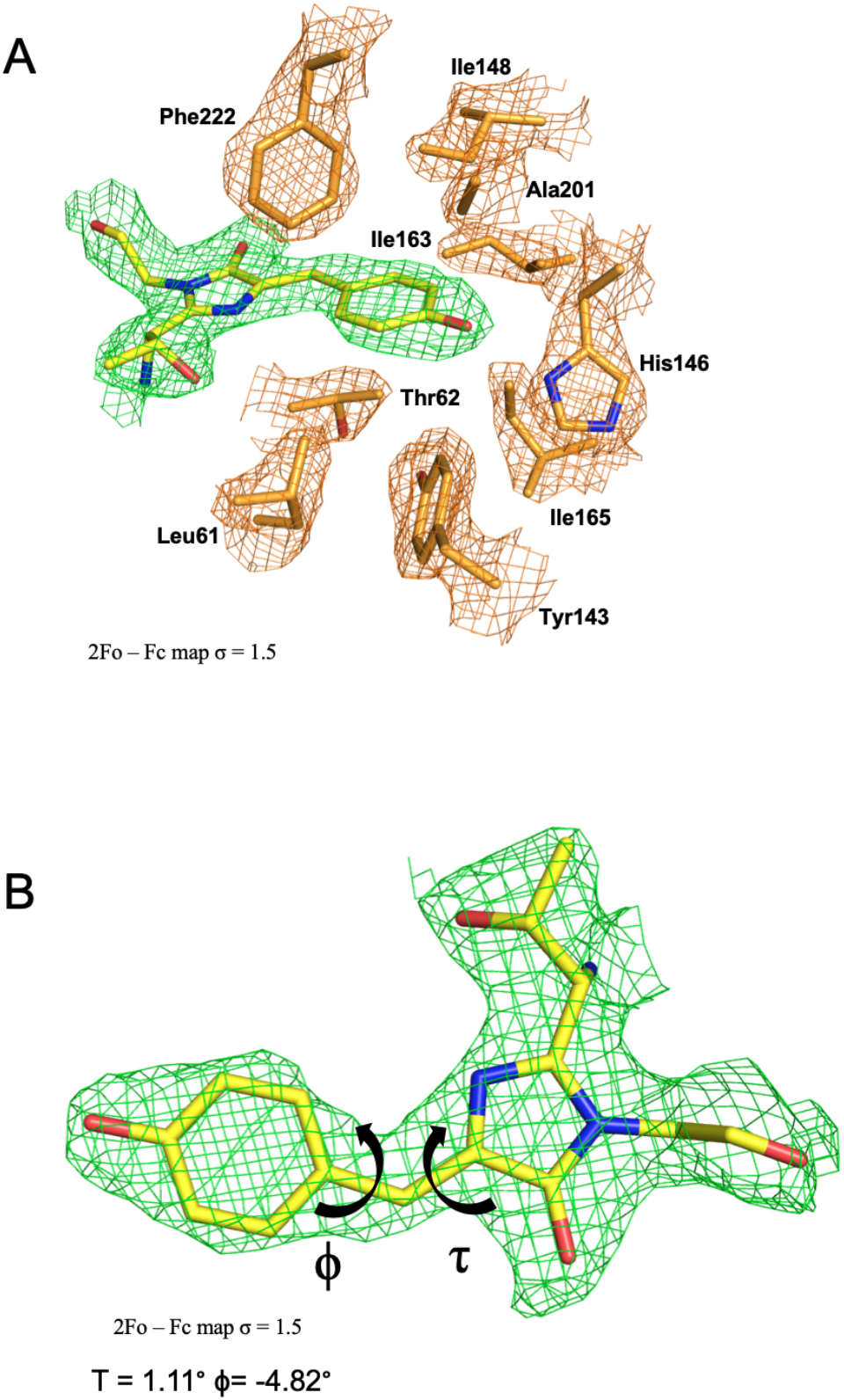
(A) Chromophore environment and (B) geometry in AausFP1.

### Unusual *Aequorea* Chromoproteins

In retrospect, the presence of green- and red-absorbing CPs in *Aequorea* species is not surprising, given the diversity of FP homologs found in other hydrozoans [21–24]. However, the properties of *Aequorea* CPs differ in unexpected ways from those previously cloned from other organisms. Every *Aequorea* CP displays a broad absorbance spectrum that lacks the well-defined sharp peak and short-wavelength shoulder typical of most FPs and CPs, suggesting that these proteins contain an unusual chromophore and/or chromophore environment. Also, none of the *Aequorea* CPs has any measurable red fluorescence emission, even on our most sensitive instruments. To our knowledge, there have been no previous reports of chromoproteins with a quantum yield of absolutely zero.

**AausFP2** has a distinctive cyan-blue pigmented appearance when expressed in *E. coli*, with a broad absorbance spectrum peaking at 610 nm. Confirmed by X-ray crystallography, AausFP2 is an obligate dimer, and sequence homology between the CPs studied suggests that they are all dimers. *A*. cf. *australis* expresses a second CP, **AausFP3**, that displays a similarly symmetrical, shoulder-less absorbance peak, but with a maximum absorbance at 590 nm.

The X-ray crystal structure of AausFP2 further revealed a chemically novel chromophore in which the side chain of a neighboring cysteine is covalently linked to the methylene bridge of a twisted GFP-like chromophore (**Fig. 7**). This amino acid, Cys62, is conserved in all *Aequorea* chromoproteins. The C62S mutant of AausFP2 appears yellow, and has a major absorbance peak characteristic of a GFP-type chromophore (data not shown), strongly suggesting that this conserved cysteine is necessary for formation of the red-shifted chromophore. The peak absorbance wavelength of alkali-denatured *Aequorea* CPs displays a 20-30 nm red-shift relative to that expected for a GFP-type chromophore [2] which is abolished by addition of β-mercaptoethanol (data not shown), providing additional evidence for the role of this unusual bond. Quantum mechanical calculations indicate that both the presence of a sulfur atom and a twisted chromophore are required to produce long-wavelength absorbance (see **Supplementary Results and Discussion**).

**Figure 7.**
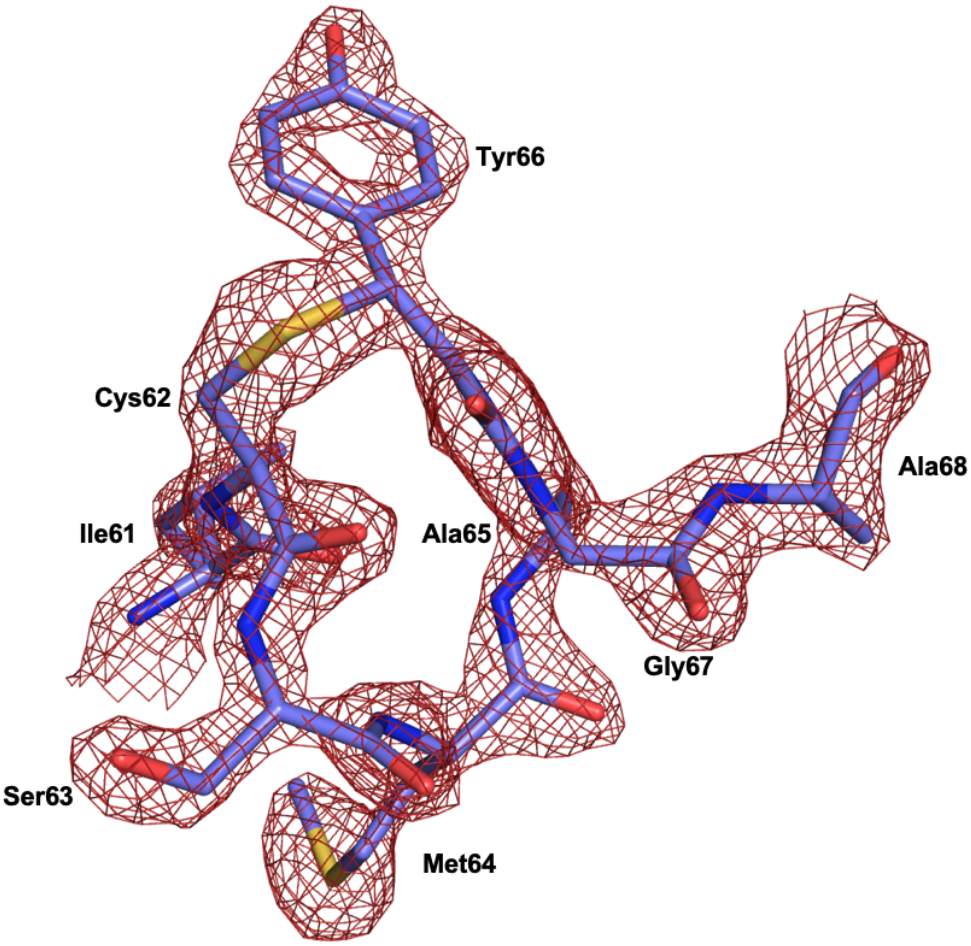
2Fo – Fc map, σ = 2.0, of the chromophore and nearby connected residues for AausFP2.

Unlike their orthologs in *A*. cf. *australis*, which mature fully to their long wavelength forms in the dark, the *A. victoria* CPs have the surprising additional requirement for blue light in order to become CPs. When expressed in total darkness, **AvicFP2** is weakly green fluorescent with spectra suggesting an avGFP-type chromophore. After blue light exposure, AvicFP2 rapidly converts (within seconds to minutes), to a purple-blue CP (peak absorbance at 588 nm) which is completely non-fluorescent. In alkali-denaturation experiments, we observed a 20 nm shift in absorbance maxima between the unconverted and converted forms of AvicFP2 (data not shown), suggesting that the Cys62 side chain becomes attached to the chromophore only *after* photoconversion. **AvicFP3** is highly homologous to AvicFP2 (96% amino acid identity, see **Fig. 4**), and is similarly green fluorescent when expressed and purified in the dark. Like AvicFP2, AvicFP3 converts to a green-absorbing CP when exposed to blue light. We suspect that AvicFP3 is so highly sensitive to blue light exposure that the act of taking an absorbance measurement is enough to partially photoconvert the protein (see **Fig. 2**).

### A Reversibly Photochromic FP

The final surprising discovery among the *Aequorea* species FPs is **AausFP4**, a very weakly fluorescent (quantum yield < 0.001) green-emitting FP with photochromic behavior strikingly similar to that of the engineered avGFP variant Dreiklang [26]. When expressed or stored in the dark, AausFP4 reaches an equilibrium state with a major absorbance peak at 338 nm, indicating that the chromophore is neutral and missing at least one double bond relative to a mature GFP-type chromophore. Upon exposure to UV light, AausFP4 rapidly and fully converts to an anionic GFP-like state with 477 nm peak absorbance.

This transformation is reversible by exposure to bright blue light or by storage in the dark. Together, these properties suggest a mechanism similar to that of Dreiklang, in which a structural water molecule can reversibly hydrate the imidazolinone ring of the chromophore in a light-dependent manner [26]. A key difference between AausFP4 and Dreiklang is the absence of a ~400 nm absorbance peak in the “on” state, accompanied by off-switching mediated by blue rather than violet light. While AausFP4 is likely to be dimeric like its closest relatives (AausFP2 and AausFP3), it may prove to be a useful starting material from which to engineer a new lineage of reversibly photoswitchable FPs. AausFP4 also represents, to our knowledge, the first naturally-occurring example of Dreiklang-type photoswitching to be discovered.

## Conclusion

We have identified several new *Aequorea* FPs with the potential to further diversify the landscape of fluorescent probes and biosensors. AausFP1, the brightest fluorescent protein currently known, will serve as the parent of an entirely new lineage of super-bright FP variants. As an apparently superior scaffold to avGFP in many ways, mAvicFP1 may be quickly adaptable to existing probes and biosensors. AausFP4 is the first natural example of Dreiklang-type photochromism, and may help generate other useful variations on this mechanism. Four highly unusual *Aequorea* chromoproteins provide truly novel engineering opportunities, including generating new far-red-emitting FPs, improved dark FRET acceptors, and photoacoustic probes, among many other potential uses.

The discovery and understanding of these new fluorescent proteins in *Aequorea* was made possible through a highly collaborative and interdisciplinary approach involving field collection work, basic molecular biology, next-gen sequencing and bioinformatics, protein engineering, microscopy, X-ray crystallography, and phylogenetics. We are optimistic that more studies with this kind of holistic approach will help elucidate many of the mysteries still hiding in the natural world. In the time that has elapsed since Shimomura’s first sampling of *Aequorea victoria* in Friday Harbor, it has become clear that there is an urgent need to explore and understand as much of the molecular biodiversity that exists in the world before many organisms go extinct or become too rare to sample.

## Supporting information

Movie 1, LifeAct

A. cf. australis native FP nucleotide sequences

A. victoria native FP nucleotide sequences

Aequorea FP peptide sequences

## Acknowledgements

We dedicate this manuscript to the memory of Dr. Roger Y. Tsien and Dr. Osamu Shimomura, whose studies on *A. victoria* and avGFP continue to inspire us and to catalyze new technologies for biological imaging. This work was also made possible by the Crystal Jelly exhibit at the Birch Aquarium at Scripps, highlighting the significance of this species in the history of biomedical research. This exhibit allows guests to visually observe the fluorescence in the jellies with the push of a button, and was the source of the *A. victoria* individual used in this work. We also wish to thank Dr. Lauren M. Barnett for aiding in the collection of *A*. cf. *australis*, Wyatt Patry (Monterey Bay Aquarium) for helping in species identification, and Dr. Ute Hochgeschwender, Dr. Stephen R. Adams, Dr. Thomas Blacker, and Dr. Robert E. Campbell for helpful feedback on the manuscript.

This work was supported by NIH R01GM109984 (NCS), and NIH R01GM121944 (NCS), NIH U01NS099709 (Hochgeschwender, Moore, and NCS), and NSF NeuroNex 1707352 (Moore). All plasmids described in this manuscript ***will be*** deposited with AddGene and crystal coordinates ***will be*** deposited with RCSB. Spectra, sequences, and photophysical characterization data ***will be*** deposited on FPbase.org. *Prior to deposit, interested non-profit researchers are encouraged to contact the corresponding author with sample and/or data requests*.

A substantial amount of additional data and discussion are included in the **Supplementary Materials** document accompanying this manuscript.

## Author Contributions

GGL performed RNA extraction, molecular cloning, protein expression and purification, optical and biochemical characterization of FPs, design and generation of mutants, synthesized data and written material provided by co-authors, and wrote the manuscript. HD, GG, and AR performed protein crystallization, X-ray crystallography, and structural analysis. DTS coordinated hydrozoan sample collection, performed morphological and molecular species identification of *A*. cf. *australis*, and reanalyzed and reconstructed FP-encoding transcripts. IN performed quantum mechanical calculations on model chromophores. DSB and TL performed microscopy and image analysis, including the OSER assay. AS provided the collection permit, supplied expertise in animal collection and care in the field, and performed optical analysis of invertebrate tissue samples. VL and JM, Birch Aquarium at Scripps, supplied cultured *A. victoria*. NCS provided funding from sponsored grants, project guidance and coordination of collaborative efforts, and performed transcriptome assemblies. HD, GG, DTS, IN, AR, DSB, TL, JM, and NCS contributed to providing material for and editing the manuscript.

## Video file descriptions

**Movie 1.** Confocal imaging of H2B-mAvicFP1 expressed in U2-OS cells. Scale bar is 10 μm. Timestamp is in hh:mm. Images were taken at 3 min intervals. Video playback is at 14 frames per second (total imaging duration 3 h 9 min).

**Movie 2.** Confocal imaging of LifeAct-mAvicFP1 expressed in U2-OS cells. Scale bar is 5 μm. Timestamp is in hh:mm. Images were taken at 4.6 s intervals. Video playback is at 100 frames per second (total imaging duration 4 h 35 min)

**Table 1.**
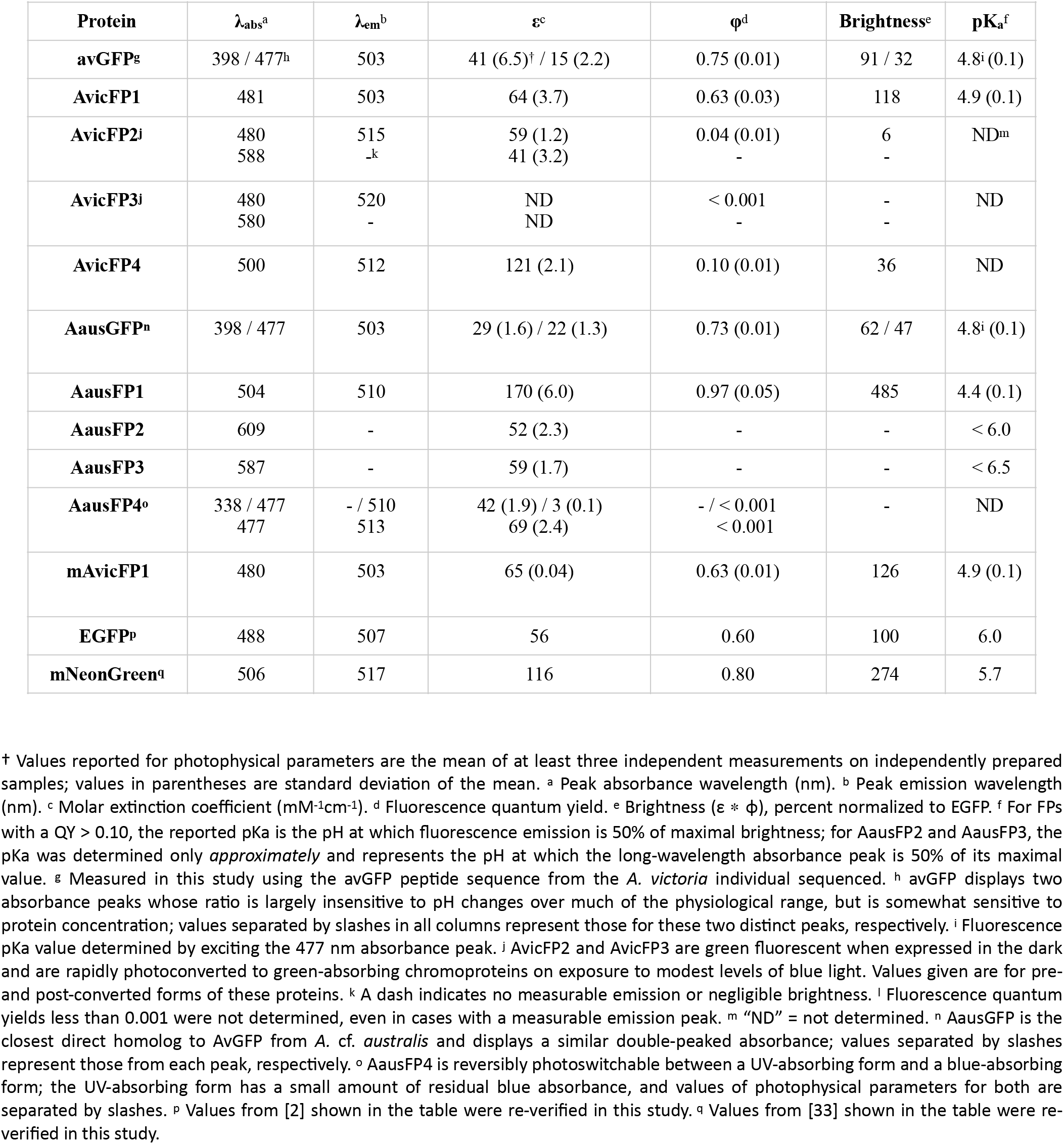
Photophysical properties of fluorescent proteins described in this study derived from *A. victoria* and *A*. cf. *australis*. The commonly used protein EGFP and the bright monomeric FP mNeonGreen are included for comparison.

## Materials and Methods

### Chemicals and other reagents

Unless otherwise noted, bacterial growth medium components were purchased from Fisher Scientific, antibiotics were purchased from Gold Biotechnology, and other chemicals were purchased from Sigma-Aldrich.

### Sample collection and RNA extraction

All scientific collection in the field was performed under permit G17/39943.1 granted to Dr. Anya Salih, Western Sydney University, by the Great Barrier Reef Marine Park Authority. A specimen of *A*. cf. *australis* was collected within the Scientific Research Zone surrounding Heron Island (Queensland, Australia) using a hand-held net and was transported back to the lab in seawater. Live samples were kept in fresh running seawater for minimal amounts of time after collection.

A single individual of *A. victoria* was obtained from the aquaculture collections of the Birch Aquarium at Scripps. The animals being kept in the exhibit tank at this time were originally obtained from the Aquarium of the Pacific (Long Beach, CA, USA), where they have been bred in captivity for many generations. Notably, the *A. victoria* are fed a diet of crustaceans, and so any hydrozoan-like FP transcripts identified must come from the jelly itself rather than from contamination of the RNA-seq library with prey-derived mRNAs.

Live samples were photographed and then anaesthetized with MgCl_2_ prior to being dissected. The bell margin, bell, and mouth were dissected separately, and total RNA was extracted using RNeasy Plus mini-prep kit (QIAGEN) following the manufacturer’s instructions. For *A*. cf. *australis*, the purified samples were combined and dried in a GenTegra RNA (GenTegra) tube for transport back to the United States. For *A. victoria*, samples from the three body regions were kept separate.

### Next-gen sequencing

Total RNA samples were used as input to generate Illumina-compatible RNA-Seq libraries at the Scripps Research Institute NGS Sequencing Core facility. Total RNA underwent polyA selection prior to Illumina TruSeq library prep. Libraries were run on one NextSeq flowcell and generated between 25 and 35 million 150bp paired-end reads per sample.

Transcriptomes for individual samples as well as the aggregate *A. victoria* transcriptome were assembled using Trinity [27,28] either on a custom workstation in the lab or using the public Galaxy bioinformatics server [29]. Read mapping was performed using bowtie2 alignment [30]and RSEM [31]for cross-sample comparison. Additional details on transcript verification are included in the **Supplementary Materials**.

### Species identification

The identity of *A*. cf. *australis* was established using phylogenetic analysis, see detailed methods and results in **Supplementary Materials**. The identity of *A. victoria* was verified by the presence of an assembled transcript encoding avGFP, as well as its well-characterized morphology.

### Cloning and mutagenesis

Candidate FP-encoding transcripts were identified by BLAST homology searching using avGFP as the query against the assembled transcriptome databases as well as intermediate assembly files created by the Trinity workflow. Searching through intermediate assembly files allowed us to identify potential alternative transcript sequences and those that were (possibly incorrectly) collapsed into single contigs by Trinity. Putative FP-encoding transcripts were validated against raw read data and reconstructed as necessary (see below for detailed methods, results, and discussion). Sequence alignments were performed using Clustal Omega [25].

For each avGFP homolog identified, the coding region was identified and a synthetic gene was designed to produce the encoded polypeptide sequence using codons optimized for both human and *E. coli* expression using an in-house BioXP3200 instrument (SGI-DNA, La Jolla, CA) or ordered as a gBlock double-stranded gene fragment (Integrated DNA Technologies, San Diego, CA). Both PCR-amplified and synthetic cDNAs contained additional nucleotides at the 5’ end (GAAAACCTGTACTTCCAGGGT) and 3’ end (CGTTTGATCCGGCTGC). Fragments encoding FPs were inserted using Gibson assembly [32] into the vector pNCST (modified from [33]) that had been PCR-amplified with the oligos pNCST-vec-F and pNCST-vec-R. The pNCST plasmid contains a synthetic promoter that drives high-level constitutive expression in most *E. coli* strains. This plasmid encodes an N-terminal 6xHis tag and linker followed by a TEV protease cleavage site just before the start codon of the inserted gene.

Site-directed mutagenesis of AvicFP1 was performed by generating two fragments of the FP coding sequence by standard PCR with Phusion polymerase (New England Biolabs) and primers as listed in **Supplementary Table S7**. Mutation(s) were placed in the overlapping sequence between fragments to facilitate Gibson assembly of full-length mutant sequences in a one-step insertion into the pNCST vector.

### Recombinant protein purification

Sequence-verified plasmids were transformed into NEB5a strain *E. coli* (New England Biolabs) (because the promoter in the pNCST vector is semi-constitutive in most strains of *E. coli*, we find it convenient to use a single strain for cloning and expression), plated on LB/agar supplemented with carbenicillin (100μg/mL), and incubated overnight at 37°C. For proteins that matured efficiently at 37°C, colonies were picked and inoculated directly into a 200mL baffled wide-mouth flask containing 50mL 2xYT broth and 100μg/mL carbenicillin, and incubated overnight at 37°C with shaking at 250 RPM. For proteins requiring multiple days at room temperature to mature, a single colony was resuspended in 10mL 2xYT medium and 100μL of this suspension was plated on five 100mm petri dishes containing LB/Agar and 100μg/mL carbenicillin. After overnight incubation at 37°C to initially establish colonies, plates were then incubated at room temperature for several days in the dark._

Bacteria containing the recombinant protein were recovered by centrifuging liquid cultures in 50mL conical tubes at 4500 × g for 10 minutes. For proteins expressed on LB/agar plates, a razor blade was used to gently glide over the surface of the agar, harvesting the colonies on the blade, and then wiped into 2ml microcentrifuge tubes and gently centrifuged to the bottom of the tube. 4mL of the lysis reagent B-PER (Thermo 78248) was added for every gram of *E. coli* pellet. Tubes were gently vortexed until the pellets were completely dissolved, taking care not to form bubbles from the detergent component of the B-PER. The resulting suspension was then incubated on a gentle rocker for 15 min and then centrifuged at >20,000 × g for 10 min to pellet insoluble debris. *Note that we find that there is a strong correlation between true protein solubility and extraction efficiency in B-PER that is not true of other extraction methods such as sonication, which can solubilize aggregated FPs more readily*.

Meanwhile, we prepared a purification column by adding 1-2mL Ni-NTA resin slurry (Expedeon) into a 15mL gravity column (Bio-Rad) allowing the storage buffer to drip through. The column was equilibrated with 10 bed volumes of wash buffer (150mM Tris pH 7.5, 300mM NaCl, 5mM imidazole) and then capped at the bottom. After centrifugation, the lysate was directly added to the prepared Ni-NTA column. The column was then capped at the top and the lysate-resin slurry was tumbled end-over-end for 30 min at 4°C. The top/bottom caps were removed and the liquid was allowed to drip through by gravity flow. The column was then washed three times with three column volumes of wash buffer. Finally, the protein was eluted from the column by gradual addition of elution buffer (50mM Tris pH 7.5, 150mM NaCl, 200mM imidazole). Clear liquid was allowed to drip through, and only the fluorescent/colorful fraction was collected.

The proteins were then concentrated further using a 3kD MWCO column (Amicon/Milipore) until the volume of protein solution was < 150μL. Meanwhile, 2x desalting columns (Pierce) were prepared for each protein by equilibrating in 50mM Tris pH 8.5, 150mM NaCl according to the manufacturer’s instructions. 150μL of protein solution was loaded onto the equilibrated desalting column and centrifuged at 1500rpm for 1 min in a microcentrifuge. The collected protein was then passed through a second equilibrated desalting column to ensure complete buffer exchange.

### Mammalian cell imaging

For experiments performed in Dr. Shaner’s lab: U2-OS cells (HTB-96, ATCC) were grown in a 35mm glass bottom dish (P35G-1.5-14-C, MatTek corporation) or on coverslips (25CIRCLE #1.5, Fisherbrand) with DMEM (105666-016, Gibco) supplemented with 10 % (v/v) FBS (10437-028, Gibco) under 5% humidified CO2 atmosphere at 37°C. Polyethylenimine (PEI) in ddH2O (1 mg/ml, pH 7.3, 23966, Polysciences) was used as the transfection reagent. The transfection mixture was prepared in Opti-MEM (31985047, Thermo Fisher Scientific) with 4.5μg PEI and 500 ng plasmid. For static images, a coverslip was placed in an AttoFLuor cell chamber (A7816, Invitrogen), and FluoroBrite DMEM (A18967-01, Gibco) was added. For time series, culture media in glass-bottom dishes was replaced with FluoroBrite DMEM (A18967-01, Gibco) supplemented with GlutaMAX (35050-061, Gibco) and 10% (v/v) FBS (10437-028, Gibco).

Confocal images and time series were acquired on a Leica TCS SP8 system using a 488nm Argon laser for excitation. The sample was placed in an incubation chamber with a controlled environment at 37°C and 5% humidified CO_2_ (OkoLab). For LifeAct-mAvicFP1, a 63x/1.40 Oil objective (HC PL APO CS2 63x/1.40 Oil, 15506350) was used with an emission bandwidth of 500 – 550 nm detected with a HyD. For time series, a bandwidth of 500 – 600 nm was used and images were acquired at 4.6 s intervals (4x line averaging, pinhole at 510 nm, 1 A.U.). For single images of H2B-mAvicFP1, CytERM-mAvicFP1, and CytERM-mGFP (Addgene 62237), a 20× 0.75NA air objective (HC PL APO 20x/0.75 CS2, 15506517) was used with an emission bandwidth of 500 – 550 nm detected with a HyD. For time series of these constructs, a bandwidth of 500 – 600 nm was used, and images were acquired at 3 min intervals (4x line averaging, pinhole at 510nm, 1 A.U.).

For experiments performed at Harvard Medical School: U2-OS cells were grown on #1.5 glass-bottomed 35mm dishes (MatTek) in McCoy’s 5A medium supplemented with GlutaMAX(ThermoFisher) and 10% fetal bovine serum (ThermoFisher) and transfected with 0.5μg pCytERM-mAvicFP1 and pCytERM-mEGFP plasmid DNA using fuGENE (Promega) 24 hours prior to imaging. Before imaging, the growth media was replaced with FluoroBrite DMEM medium supplemented with 5% FBS (ThermoFisher). Image acquisition was performed in a full environmental enclosure (37°C, 5% CO2; Okolab) on a Nikon Ti-E microscope with Perfect Focus System, a Spectral Borealis-modified spinning disc confocal (Yokogawa X1), and an Orca Flash v3 sCMOS camera (Hamamatsu). OSER data was acquired with a 40X Plan Fluor 1.3 NA objective lens and live time-lapse imaging was acquired with a 100x Plan Apo VC 1.4 NA objective (162nm and 65nm pixel size respectively). Green fluorescence was excited with a 491nm solid state laser (Cobolt) and a Di01-T405/488/568/647 (Semrock) dichroic; emission was selected with an ET525/50m filter (Chroma). Hardware was controlled with MetaMorph (v7.8.13). For time-lapse experiments, single-plane images were acquired every second. For OSER acquisition, a uniform grid of images were acquired covering the entire coverslip.

Images are processed with help of Fiji [34]. OSER assay analysis was conducted as previously described [19].

### Quantum yield and extinction coefficient determination

Purified green-emitting FPs were characterized as previously described [35] to determine quantum yield. Briefly, FPs that had been buffer-exchanged into 50mM Tris-HCl pH 8.5, 150mM NaCl were diluted into the same buffer until the baselined peak absorbance was ≤ 0.05 as measured by a UV-2700 UV-Vis spectrophotometer (Shimadzu). Sample and standard (fluorescein in 0.1M NaOH, quantum yield 0.95 [36]) absorbance was matched within 10% at 480nm, the excitation wavelength used for fluorescence emission spectra. Immediately after measuring the absorbance spectrum, the cuvette containing the sample was transferred to a Fluorolog-3 fluorimeter (Jobin Yvon) and the emission spectrum was taken from 460nm to 700nm in 1nm steps, with excitation at 480nm and a slit width of 2nm for both excitation and emission. Emission spectra were interpolated under the region in which scattered excitation light bleeds through into the emission path. Quantum yield was calculated by dividing the area under the sample emission curve by its absorbance at 480nm and dividing by the same ratio for the standard, then multiplying by 0.95, the quantum yield of the standard.

Extinction coefficients for all FPs and CPs in this study were measured using alkali denaturation (addition of 2M NaOH to the FP sample to a final concentration of 1M NaOH) as previously described [35] with the following modifications: (1) In order to avoid calculating erroneously large values of FP extinction coefficients from alkali denaturation measurements, several absorbance spectra were taken for each sample. Beginning immediately after addition of NaOH, multiple absorbance spectra were taken over several minutes to determine both the point at which the protein was fully denatured and the point at which it reached maximum absorbance at ~447nm. The *maximum* measured value of the peak absorbance of fully denatured protein was used in extinction coefficient calculations. (2) For chromoproteins containing the novel cysteine-linked chromophore, the peak absorbance of alkali-denatured protein is red-shifted 20-30 nm relative to the known 447nm peak of GFP-type chromophores [2]. Because the extinction coefficient of this species is unknown, we also measured absorbance spectra for alkali-denatured CPs with the addition of 1mM β-mercaptoethanol, which is expected to break the bond between the sulfur atom of the cysteine side chain and the methylene bridge of the chromophore, producing a GFP-type denatured chromophore. The maximum absorbance value of reduced, denatured chromophore was used in calculation of the extinction coefficient, which should be considered an estimate for *Aequorea* CPs pending much deeper investigation into the biochemical properties of their unique chromophore.

### pK_a_ determination

Purified proteins were concentrated and desalted as described above into 20mM Tris-HCl pH 8. A solution of 50mM Tris-HCl, 50mM citric acid, 50mM glycine, and 150mM NaCl (final concentrations after pH adjustment) was prepared and split into two master stocks that were adjusted to pH 3 and pH 12 with HCl and NaOH, respectively. These stocks were then used to prepare buffers at pH 3, 4, 5, 6, 6.5, 7, 7.25, 7,5, 7,75, 8, 9, 10, 11, and 12 by mixing at different ratios. Each sample was then diluted (2μL sample + 198μL buffer) into each pH buffer and its emission or absorbance was measured using an Infinite M1000Pro (TECAN) plate reader. The pK_a_ was determined by interpolating the pH value at which the fluorescence or absorbance value was 50% of its maximum.

### Protein crystallogenesis

AausFP1 and Aaus FP2 were first expressed and purified as aforementioned. The His-tag was cleaved off using either TEV for AausFP1 (1/100 protease/protein ratio, overnight incubation at room temperature) or proteinase K for AausFP2 (1/50 protease/protein ratio, 1 h incubation at room temperature). The protein solution was run through an additional His-Trap column to remove cleaved tag and uncleaved protein. A final purification step consisted of a gel filtration column (Superdex 75-10/300 GL (GE Healthcare, Chicago). Fractions were analyzed using 15% SDS-PAGE gels, pooled and concentrated to 40 and 51 mg/mL for AausFP1 and AausFP2, respectively using an Amicon Ultra centrifugal filter with a molecular weight cutoff of 30 kDa (Merck, Darmstadt). Initial crystallization hits were obtained using the HTX lab platform of the EMBL Grenoble outstation, and then manually optimized. AausFP1 was crystallized with the hanging drop method using 0.7-1.3 M trisodium citrate, 0.2 M sodium chloride in 0.1M Tris buffer pH 6.5 – 8.0. AausFP2 was crystallized with the hanging drop method using 14-24% PEG 3350 trisodium citrate, 0.2 M sodium chloride in 0.1M HEPES buffer pH 7.3 – 8.2.

### Diffraction data collection

Diffraction data for AausFP1 were collected on beamline BL13-XALOC at the Spanish Synchrotron in Barcelona (Spain) [37] from a crystal flash-cooled at 100K in its mother liquor supplemented with 20% (v/v) of glycerol for cryoprotection. Diffraction data for AausFP2 were collected on beamline ID30B of the European Synchrotron Radiation Facility in Grenoble (France) [38] from a crystal flash-cooled at 100 K without addition of any cryoprotectant. Diffraction data were integrated and reduced using *XDS* and *XSCALE* [39]. Data collection and reduction statistics are given in **Supplementary Table S2**.

### Structure determination

A BLAST search (https://blast.ncbi.nlm.nih.gov/) identified the fluorescent protein phiYFPv from the jellyfish genus *Phialidium* as the closest homologue of both AausFP1 and AausFP2 (sequence identities of 61% and 50%, respectively) with a known structure (PDB entry code 4HE4, [40]). The structures of AausFP1 and AausFP2 were solved by the molecular replacement method using the 4HE4 coordinates as a search model with the program *PHASER* [41]. The model was progressively and interactively modified in COOT [42] and refined with REFMAC5 [43]. The asymmetric units contain 4 molecules for AausFP1 and 1 molecule for AausFP2. Analysis of the interaction interfaces with PISA [44] strongly suggest that the AausFP1 tetramer consists of a dimer of a physiological dimer (interface areas of 1210 Å^2^ vs. 360 Å^2^) for the third and fourth largest areas) while the AausFP2 monomer forms a physiological dimer with a symmetry-related molecule (interface area of 1290 Å^2^ *vs*. 540 Å^2^ for the second largest area). Structure refinement statistics are given in Table 3.

### Calculation of AausFP2 absorption maxima

Eight models of the minimal part of the chromophore were constructed, modelling only the two conjugated cycles of the chromophore. H atoms replaced in all models the two alpha carbone atoms linking the chromophore to the rest of the protein. 3D coordinates for all heavy atoms of the chromophore were taken from the crystallographic structures without optimization leading to two groups of models, the one with the conformation of the EGFP structure and the one with the conformation of the AausFP2 structure. The corresponding sets of models were labeled EGFP or AausFP2. The main difference between the two sets of models is the dihedral angle between the two cycles, that is −2° (almost planar) for EGFP and −53° (twisted) for AausFP2.

In each set of models, the phenol moiety was presented in its protonated form (neutral chromophore) or phenolate form (anionic chromophore). Moreover, in the AausFP2, the carbon between the two cycles of chromophore is linked to a protein’s cysteine through a thioether bond, whereas this carbon is simply protonated in the case of EGFP. Therefore, in the models, this carbon was linked either to a mercapto group (-SH) or simply protonated. Structures were protonated and the position of H atoms were optimized at the B3LYP/6-31+g(d,p) level of theory with Gaussian G09 program.

