## Supplementary Materials for "*Aequorea victoria’s* secrets"

### Supplementary Material

Figures and tables included in this supplementary material are numbered **S1, S2,** etc. All other figure and table references below are to the main text of the manuscript. Citation numbers refer to the **Supplementary References** list at the end of this document.

#### Supplementary Results and Discussion

##### Collection and identification of *Aequorea* cf. *australis* and *Aequorea victoria*

The Great Barrier Reef *Aequorea* specimen (**Fig. 1C, D**) was collected by hand net while blue-water snorkeling at -23.474046, 151.975436. The morphology of the animal was consistent with *A. macrodactyla* specimens collected over the course of several decades [1-3]. Molecular analysis of the 16S, 18S, and COI loci were inconclusive in identifying the animal to a species level, however the COI sequence was similar to that of *A. australis* (see **Morphological Identification** and **Molecular Identification**, below). Thus, we refer to this specimen as *Aequorea* cf. *australis* to indicate the uncertainty in its true species identity. The *A. victoria* specimen (**Fig. 1A, B**) was part of the dedicated Crystal Jelly exhibit at the Birch Aquarium at Scripps, in which the animals are regularly exposed to violet light and fed a diet of small crustaceans. The species identity of this sample was verified by the presence of avGFP (see below) as well as the 18S sequence, which matched most closely to that of *A. victoria* (data not shown).

##### Identification and characterization of *Aequorea*-derived fluorescent proteins

We performed mRNA-Seq of *A. victoria* and *A.* cf. *australis* followed by *de novo* transcriptome assembly and homology searches to identify contigs encoding avGFP homologs. In total, we identified five FP-encoding transcripts in *A. victoria* and five in *A.* cf. *australis.* Amongst these proteins, we found avGFP and close homologs as well as alternative green-emitting FPs with low homology to avGFP. We also discovered a surprising variety of chromoproteins—FP homologs with strong absorbance but very low emission quantum yield that act as pigments of unknown function. Optical characterization of recombinant proteins revealed several unexpected properties among the FP homologs identified in *A. victoria* and *A.* cf. *australis,* including two photoconvertible chromoproteins in *A. victoria*, a highly red-shifted chromoprotein in *A.* cf. *australis,* and a very bright green FP in *A.* cf. *australis.* Below, we describe the most notable properties of each FP. White light and fluorescence images of the full set of purified proteins are shown in **Fig. 2**, and the absorbance and fluorescence emission spectra are shown in **Fig. 3.** The photophysical properties of all FPs described here are summarized in **Table 1**, and transcript abundances for each FP in the sampled tissues are shown in **Table S8**.

***Aequorea victoria:***

**avGFP**

As one might expect given the easily visible green fluorescence of the animal, and the historical success in cloning despite much less advanced technology, we found that transcripts encoding avGFP were the most highly represented of the FP homologs in the *A. victoria* transcriptome, with highest expression in the bell margin and minor presence in other tissues. Interestingly, despite the bright green fluorescence around the bell margin, we found that this transcript was still represented at relatively low levels (23 transcripts per million (TPM)). We hypothesize that avGFP may have a long half-life *in vivo* and thus require relatively little active transcription to maintain fluorescence in the animal.

The consensus contig for avGFP in the individual *A. victoria* sequenced in this study encodes a protein that differs by one or two conservative amino acid substitutions relative to the originally published avGFP [4]. In our hands, the optical and biochemical properties of the protein remain unchanged relative to its original description. These minor sequence differences between the historically known avGFP and the transcript identified here could be explained by population segregation between Friday Harbor (Shimomura's original *A. victoria* collection location) and central California, where the ancestors of our specimen were originally collected by the Aquarium of the Pacific, which then supplied cultured animals (an unknown number of generations later) to the Birch Aquarium at Scripps, the source of the sequenced individual.

**AvicFP1**

The first surprise among the newly discovered *A. victoria* FPs was an FP with relatively high homology to avGFP (81% amino acid sequence identity, see **Fig. 4**). This transcript was represented at much lower levels in our RNA-Seq data, and was present only in the "body" and mouth of the animal (~0.06 and 0.38 TPM, respectively). Unlike avGFP, AvicFP1's absorbance spectrum indicates that it contains an avGFP-type chromophore in the fully anionic state, with a peak at 481 nm and a fluorescence pKa of 4.9, substantially lower than that of EGFP . We attribute this difference largely to the first position in the chromophore tripeptide, which is a cysteine in AvicFP1 rather than the serine present in avGFP. The mutation S65T in avGFP is among the most critical early mutations introduced to generate an all-anionic chromophore, and S65C has been reported as well [5,6].

Because we generated the *E. coli* expression plasmids using synthetic genes, some clones also contained additional mutations derived from errors in the oligonucleotides used for gene assembly. We identified one colony among these initial clones that produced a much larger proportion of mature FP at 37ºC. The sequence revealed this clone contained the mutation F64L, another of the early folding mutations identified for avGFP [7]. Unlike avGFP, which required a large number of additional mutations to become truly useful, AvicFP1-F64L already appears to fold and mature nearly optimally under standard mammalian tissue culture conditions with only this single point mutation, which maintains both the brightness and low pKa of the wild-type FP.

A careful examination of the sequence alignment between AvicFP1 and avGFP revealed that essentially all of the side chains that participate in the weak dimer interface of avGFP are conserved in AvicFP1. Based on this observation, we hypothesized that mutations sufficient to monomerize avGFP variants (i.e. A206K [8]) would also produce a monomeric variant of AvicFP1. Using the organized smooth endoplasmic reticulum (OSER) assay to test for oligomeric behavior in cells [9], we found that the mutant AvicFP1-F64L/A206K displays monomeric behavior equivalent to mEGFP, widely considered the "gold standard" of monomeric FPs [9] (OSER data are summarized in **Table S3**). Fusions to LifeAct and H2B displayed the expected localization (**Fig. 5, Movie 1,** and **Movie 2**), and did not appear to interfere with mitosis or cell growth based on qualitative observations. We therefore designated this protein "monomeric *A. victoria* fluorescent protein 1," or **mAvicFP1**.

By all metrics we have evaluated, mAvicFP1, which contains ***only two mutations*** relative to wild-type AvicFP1, is superior to mEGFP in our hands. Its quantum yield and extinction coefficient are higher, and it can tolerate a wider pH range than mEGFP, making it potentially useful for imaging fusions localized to acidic compartments. We did not explicitly measure the photostability of mAvicFP1, but based on qualitative observations, it appears similar to that of mEGFP. We additionally generated blue (Y66H) and yellow (T203Y) variants of AvicFP1 (data not shown), indicating that this protein is likely to be amenable to additional engineering and optimization. Because this study was focused primarily on identifying and characterizing all of the newly discovered FPs in *Aequorea* species, we did not extensively test mAvicFP1 as a fusion tag beyond our initial experiments, and it is likely that mAvicFP1 can be further improved by directed evolution. However, we soon refocused our efforts on monomerizing **AausFP1** (see below), a much brighter green-emitting FP with much lower homology to avGFP.

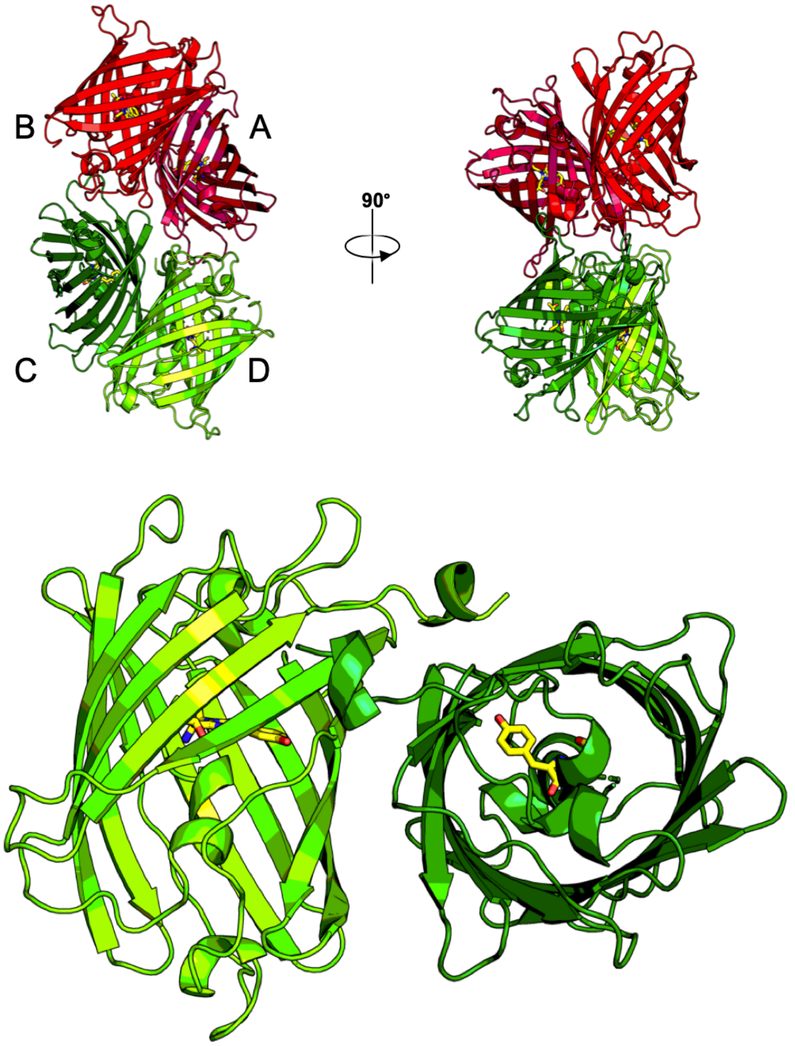

**Figure S1.** Cartoon diagram of the AausFP1 unit cell and likely physiological dimer. The four protein subunits in the unit cell are labeled A through D.

**AvicFP2** and **AvicFP3**

The first chromoprotein (CP) characterized from *A. victoria* matures to a purple-blue color that appears to be completely non-fluorescent, even when measured on our most sensitive instruments. Interestingly however, the chromophore requires blue light to mature to this long-wavelength-absorbing state; when expressed and purified in total darkness, AvicFP2 is weakly green fluorescent with excitation and emission spectra suggesting an anionic avGFP-type chromophore. Exposure to a moderate intensity of blue light induces rapid conversion to the mature chromoprotein state within seconds to minutes. Like all other *Aequorea* species CPs discovered in this study, AvicFP2 contains a conserved cysteine residue that is most likely responsible for forming a previously undescribed chromophore structure in the fully mature state (see detailed discussion below).

AvicFP3 is a second chromoprotein that is highly homologous to AvicFP2 (96% amino acid identity, see **Fig. 4**). Like AvicFP2, AvicFP3 is green fluorescent when expressed and purified in the dark, and converts to a green-absorbing chromoprotein when exposed to blue light. AvicFP3 qualitatively appears to be more light-sensitive in its immature state than AvicFP2, making measurement of the fully immature protein nearly impossible in our hands. Alternatively, AvicFP3 may be capable of partially maturing in the dark under some conditions. Both AvicFP2 and AvicFP3 transcripts were only found in the "body" of the animal, and were expressed at low levels (~0.38 and ~0.09 TPM, respectively).

**AvicFP4**
One transcript in the initial *A. victoria* transcriptome assemblies appeared to encode an additional incomplete GFP homolog, however the transcript appeared to be misassembled upon inspection with mapped reads. We therefore went back and used only the reads mapping to this partial GFP transcript as the input for a new *de novo* assembly and found that one contig in this assembly encoded a full-length ortholog to AausFP1 (discussed below). This transcript is present in all sampled tissues at moderate levels (between 1.67 and 5.11 TPM). AvicFP4 has low sequence identity to avGFP (52% amino acid identity). When expressed in *E. coli* at 37°C, AausFP4 folds and matures efficiently into a highly soluble green-emitting FP with narrow peaks and a short stokes shift. While it has a relatively high extinction coefficient, its quantum yield is much lower than is typical of wild-type green-emitting FPs. Thus, while this protein may provide many interesting insights into determinants of quantum yield and spectral shape, we did not characterize it beyond measuring its core optical properties.

***Aequorea* cf. *australis:***
**AausGFP**
*A.* cf. *australis* expresses one fluorescent protein that appears to be an ortholog to avGFP (81% amino acid identity). AausGFP has optical and biochemical properties very similar to those of wild-type avGFP, suggesting that it may also be functionally equivalent in this species. However, AausGFP was expressed at relatively low levels (9 TPM) compared with other FP homologs in the same animal. Like avGFP, enhanced green, yellow, cyan, and blue variants of AausGFP can be produced with similar amino acid substitutions to those used to make the avGFP color variants (data not shown). AausGFP can likely be monomerized similarly to avGFP, given the high degree of sequence conservation between the proteins. However, since its properties are neither novel nor superior to existing FPs, we did not explore this protein beyond these initial characterization and mutagenesis experiments.

**AausFP1**
Among the many surprises amongst the *Aequorea* species transcriptomes studied here, we discovered a second green-emitting FP in *A.* cf. *australis* with very low homology to avGFP (53% amino acid identity). AausFP1 was expressed at very low levels relative to other FPs in the individual sequenced (0.36 TPM), and would be unlikely to appear in cDNA expression-cloning libraries, thus making the transcriptomic approach used in this study one of the few approaches capable of elucidating the existence of this FP. In fact, only 43 reads out of ~25M reads mapped to the AausFP1 transcript (see **Verifying Newly Identified Transcripts** and **Fig. S-F,** below). Despite low expression in its native context, wild-type AausFP1 expresses and folds very efficiently in *E. coli* at 37ºC without any modifications.

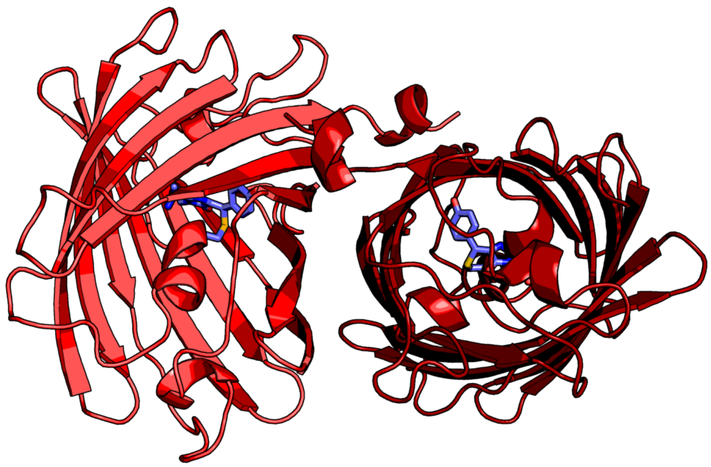

**Figure S2.** Cartoon diagram of the probable AausFP2 physiological dimer.

Most importantly, AausFP1 displays truly remarkable optical properties: narrow excitation and emission peaks, nearly perfect QY (close to 1.0), and the highest extinction coefficient yet measured for any fluorescent protein (170,000 $M^{-1}cm^{-1}$). Together, these properties make AausFP1 the brightest fluorescent protein currently known in the avGFP superfamily, including engineered variants. The emission peak of

AausFP1 has a full width at half maximum (FWHM) of 19nm, compared to 40nm for fluorescein, 32nm for EGFP, and 28nm for mNeonGreen (calculated from spectra taken in this work). Such a narrow emission spectrum is highly desirable for multi-color fluorescence imaging because it minimizes bleed-through to long wavelengths while allowing efficient collection of emission light from a narrow wavelength band. However, like many wild-type hydrozoan fluorescent proteins, AausFP1 is an obligate dimer, as verified by X-ray crystallography.

**AausFP2** and **AausFP3**

The most highly expressed FP homolog in *A.* cf. *australis* is a chromoprotein with a cyan-blue pigmented appearance when expressed in *E. coli* (see **Fig. 2**). AausFP2 possesses an unusually-shaped, broad absorbance spectrum peaking at 610 nm. The apparent absence of a classical sharp peak with a short-wavelength shoulder indicates that the chromophore and/or chromophore environment of this protein must also be highly unusual. Indeed, this is what we discovered in the solved crystal structure (detailed discussion below). Like the *A. victoria* chromoproteins, AausFP2 has no measurable fluorescence emission, even on our most sensitive instruments. Like all *Aequorea* species chromoproteins, AausFP2 appears to be dimeric when run on a non-denaturing SDS-PAGE gel (this oligomerization state was also confirmed by X-ray crystallography, see below). AausFP2 was represented far more highly than any other FP found in this study, at over 2,000 TPM.

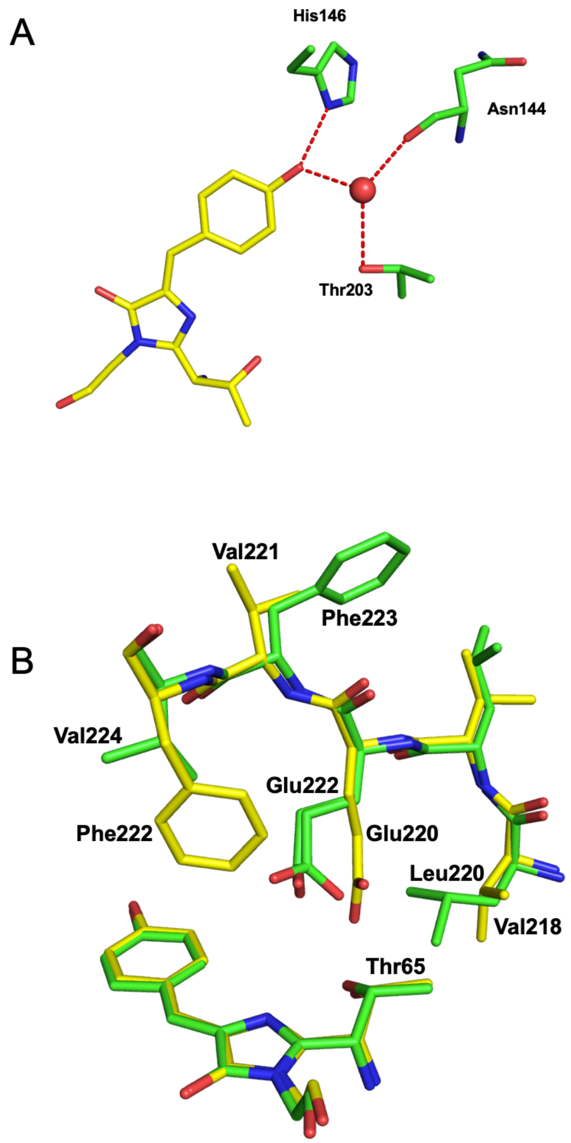

**Figure S3.** Hydrogen bonds stabilizing the AausFP1 chromophore phenolate (A) and a comparison between AausFP1 (yellow) and EGFP (green) positioning of residues near the chromophore.

*A.* cf. *australis* expresses a second chromoprotein, AausFP3, similar to AausFP2, but blue-shifted approximately 20 nm relative to its closest homolog. AausFP3 displays a similarly symmetrical, shoulder-less absorbance peak, but with a maximum at 590 nm. The transcript encoding AausFP3 is present at lower levels than that of AausFP2 (~18 TPM). Unlike the *A. victoria* chromoproteins, AausFP2 and AausFP3 do not require light for maturation, but instead form their mature blue-pigmented chromophores equally well in the dark. The potential biological functions of these chromoproteins, as well as those from *A. victoria*, are currently unknown.

**AausFP4**

The final surprising discovery among the *Aequorea* species FPs is AausFP4, a very weakly green-emitting FP (quantum yield $< 0.001$) with photochromic behavior strikingly similar to that of the engineered avGFP variant Dreiklang [10]. AausFP4 is expressed at moderate levels in the individual sequenced in this study (~87 TPM). When expressed or stored in the dark, AausFP4 reaches an equilibrium state with a major absorbance peak at 338 nm, indicating that the chromophore is neutral and missing at least one double bond relative to a mature GFP-type chromophore. Upon exposure to UV light,

AausFP4 rapidly converts entirely to an anionic GFP-like state with 477 nm peak absorbance. This transformation is reversible by exposure to bright blue light or by storage in the dark. Together, these properties suggest a mechanism similar to that of Dreiklang, in which a structural water molecule can reversibly hydrate the imidazolinone ring of the chromophore in a light-dependent manner [10]. A key difference between AausFP4 and Dreiklang is the absence of a ~400 nm absorbance peak in the "on" state, and off-switching mediated by blue rather than violet light. While AausFP4 is likely to be dimeric like its closest relatives (AausFP2 and AausFP3), it may prove to be a useful starting material from which to engineer a new lineage of reversibly photoswitchable FPs. To our knowledge, AausFP4 is also represents the first naturally-occurring example of Dreiklang-type photoswitching to be discovered.

#### Novel FP structural features

The highly unusual optical properties of AausFP1 and AausFP2 made them attractive targets for structural characterization. We obtained X-ray diffraction from crystals of these two FPs and solved their structures at 2.47 and 2.06 Å resolution, respectively. Data collection and reduction statistics are given in **Table S2.**

**Oligomerization state analysis.** While both crystal forms contain two different asymmetric unit contents with 4 monomers for AausFP1 (**Fig. 6**) and 1 monomer for AusFP2, both structures strongly suggest that the two proteins are physiologically organized as dimers (see Methods section) (**Fig. S1**, **Fig. S2**). The dimer interfaces of these two proteins are highly similar to each other (**Fig.s S13** and **S14**), but, surprisingly, are completely different from that of avGFP [11]. The analysis of residue interactions at these interfaces has provided crucial information for successfully engineering monomeric variants of these proteins (to be published separately).

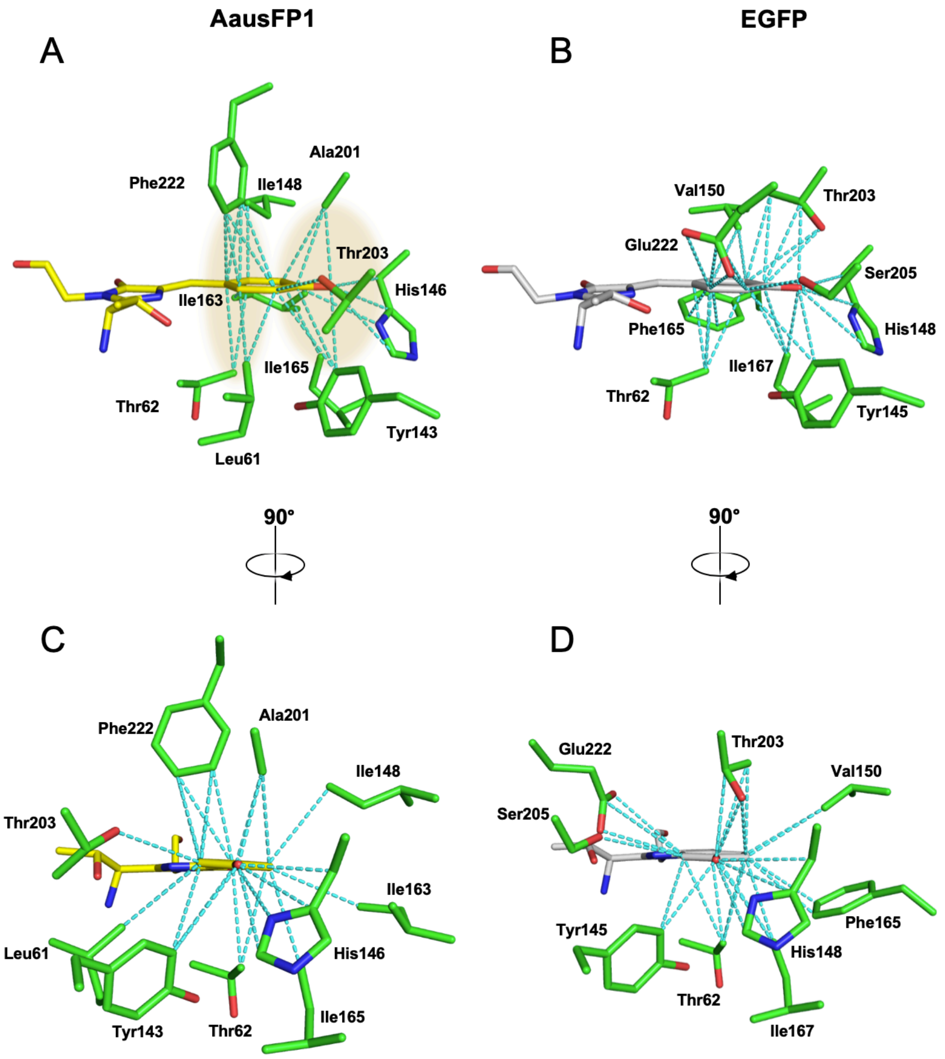

**Figure S4.** Two side-on views of van der Waals interactions in the AausFP1 and EGFP chromophore environments. AausFP1 (A, C) and EGFP (B, D) are viewed from the "right side" of the chromophore (A, B) and from the angle looking head-on at the chromophore phenolate.

**Structural analysis of the AausFP1 chromophore.** Particularly high values of the fluorescence quantum yield and molar extinction coefficient in an FP suggest the presence of an unusual chromophore conformation and/or environment. At first glance at the AausFP1 chromophore, its almost perfect planarity (**Fig. 6**) and its stabilization via hydrogen bonds of the (deprotonated) phenolate oxygen to a neighboring histidine residue and a stable water molecule (**Fig. S3a**) resemble those of other common fluorescent proteins such as EGFP ("Enhanced" avGFP, an early and still commonly-used derivative) [7].

However, a closer comparison with the chromophore environment of EGFP, which contains a chemically identical chromophore derived from a Thr-Tyr-Gly tripeptide (**Fig. S3b**), indicates that the glutamate residue conserved almost universally in known FPs (Glu220) is translated towards the chromophore residues (n-1) (Thr65) and (n+1) (Gly67), which displaces a conserved negative charge interacting with the delocalized electron cloud of the chromophore. It is unclear what effect this change may have on the quantum yield or extinction coefficient of the chromophore or its spectral shape.

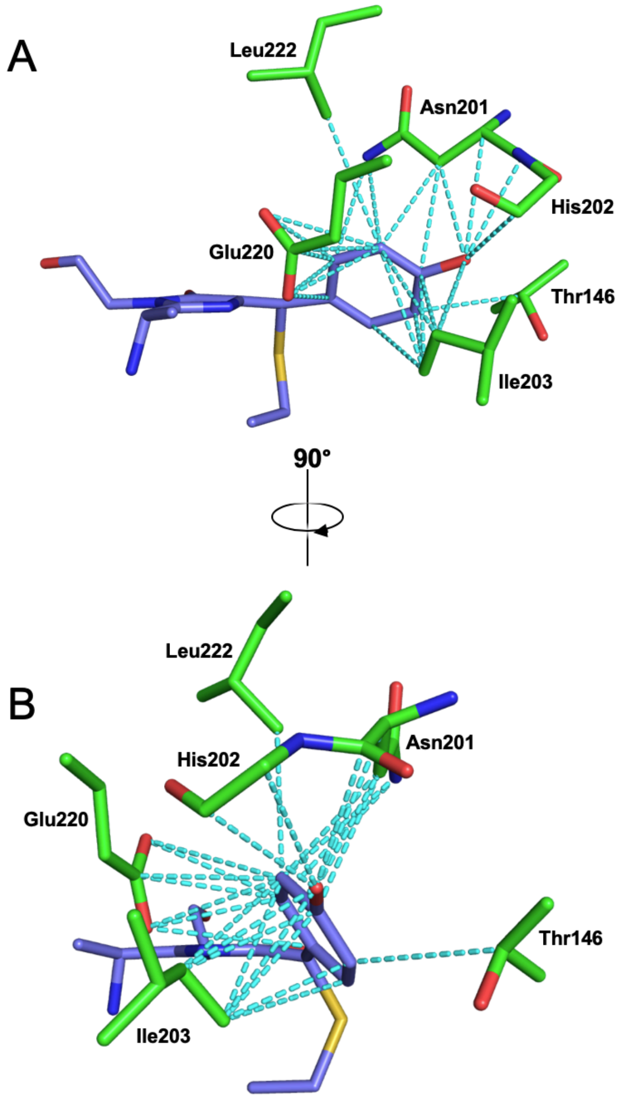

**Figure S5.** Two side-on views of van der Waals and other interactions in the AausFP2 chromophore environment

The most striking features of the AausFP1 chromophore environment are revealed by analysis of the interactions surrounding the phenolate ring (**Fig. 6a** and **Fig. S4**), which are almost exclusively van der Waals interactions. In EGFP, these interactions weakly engage residue atoms less than 4.0 Å from the phenolate ring (**Fig. S3b**). Interestingly the 5 shortest interatomic distances for van der Waals interactions with the EGFP chromophore (≤ 3.5 Å) all include an oxygen (polar) atom (see **Table S4**). The EGFP chromophore environment also features an angular sector around the phenolate ring that is entirely deprived of interactions (**Fig. S4d**), potentially leaving room for the chromophore to deform when relaxing from the excited state, and thus releasing its energy via a non-radiative pathway.

Unlike EGFP, there are many residues in AausFP1 that interact with the phenolate ring via (hydrophobic) carbon atoms. These interactions can be visualized as two larger "rings": one around atoms $C_\gamma$, $C_{\delta 1}$ and $C_{\delta 2}$ and one around $C_{\varepsilon 1}$, $C_{\varepsilon 2}$, $C_\zeta$ and $O_\eta$ (beige areas on **Fig. S4a**). Among the 6 closest interactions (≤ 3.5 Å), 3 include an oxygen atom and 3 include only carbon atoms (see **Table S5**). The side-on view of the chromophore (**Fig. S4c**) reveals that the asymmetry of interactions present in EGFP is abolished in AausFP1, and that the phenolate ring is perfectly locked in place by all interacting residues, essentially restricting any potential deformation around the methylene bridge. These substantial differences provide a sound explanation for the very high quantum yield and unusually narrow excitation and emission peaks of AausFP1 relative to EGFP.

**Structural analysis of the AausFP2 chromophore.** Unlike the AausFP1 chromophore, the configuration and conformation of the AausFP2 chromophore are unlike those of any previously described fluorescent protein homolog. The residue one turn away from the chromophore on the N-terminal side of the chromophore-bearing helix is a cysteine (Cys62) which is engaged in a covalent bond with the central carbon ($C_\beta$) of the methylene bridge of the chromophore (**Fig. S5**). This covalent bond is equivalent to the substitution of a hydrogen by a sulfur atom, preserving the delocalization of the electron cloud and the visible light absorption properties of the chromophore. In addition, the residues surrounding the AausFP2 chromophore provide an asymmetric environment that results in a further twisting of the chromophore thanks to a van der Waals 'clamp' formed by the residues Asn201, Ile203, and Glu220 (**Fig. S5**). We verified the role of Cys62 by examining the C62S mutant of AausFP2, and as predicted, the mutant was unable to form a red-absorbing chromophore and instead absorbed strongly in the blue region of the visible spectrum (data not shown).

**Modeling of the AausFP2 chromophore absorption properties.** The effects of the sulfur substitution at the Cβ position in an EGFP-like chromophore were modeled with quantum mechanical calculations and appear to be, in combination with the twisting of the two rings, the most probable explanation for the large red-shift in absorbance of AausFP2 relative to EGFP. Calculations of the energetic vertical transition between the ground state and the first six excited states were done on models of the chromophore constructed from the crystallographic structures of avGFP-F64L/S65T (Protein Data Bank entry code 2Y0G, [12]) and the structure of AausFP2 reported here. Calculations of the vertical transition on small models reproduced the large red-shift of the absorption maximum of AausFP2 relative to EGFP, and shows that this is due to both the presence of a sulfur atom in the chromophore and to the twisting of the structure due to the covalent bond to the cysteine.

Calculations were performed on a set of a total of 8 models in order to analyze the influence on the absorption wavelength of the conformations, the protonation state and the presence of the mercapto group. The models (name and formula) are gathered in **Fig. S6**. For each model, the time-dependent density functional theory (TD-DFT) calculations of the vertical transitions were conducted for the first 6 singlet excited states at the B3LYP/6-31+G(d,p) level [13,14]. The wavelengths of the most intense transition (bigger oscillator strength) are reported in **Table S1**. The main contribution of the transition has a HOMO-LUMO character.

The models that best represent the chromophore in AausFP2 are the protonated and deprotonated forms Anionic-$SH_{AausFP2}$ and Neutral-$SH_{AausFP2}$. The corresponding absorption maximum wavelength (640 and 690 nm) represent large red shifts compared to those of the anionic form of the chromophore in EGFP (Anionic-$H_{EGFP}$) (470 nm). As models do not take into account the environment, the present calculations only give trends on the position of absorption maxima and not their precise values.

The presence of the cysteine in the structure of the AausFP2 protein can influence the vertical transition in two ways: (1) the presence of the electronic density of the sulfur atom in the electronic states and/or (2) the steric constraint on the structure of the chromophore. In order to decipher the two influences, we can first compare the substitution of the -H by -SH group on the predicted peak absorbance wavelength. For the planar chromophore structure (derived from EGFP), the substitution has ***no influence*** on the wavelength of either the neutral or anionic forms of the chromophore. The transition is, however, less intense (smaller $f$ values). For the twisted chromophore structure (derived from AausFP2), the substitution of -SH by -H produces a blue shift of the wavelength of 140 and 260 nm for anionic and neutral forms, respectively. From this, we conclude that the substitution of H by SH has an influence only when the structure is twisted.

| Neutral-$H_{EGFP}$ | Anionic-$H_{EGFP}$ | Neutral-$SH_{EGFP}$ | Anionic-$SH_{EGFP}$ |
|---|---|---|---|
| Neutral-$H_{AausFP2}$ | Anionic-$H_{AausFP2}$ | Neutral-$SH_{AausFP2}$ | Anionic-$SH_{AausFP2}$ |
| 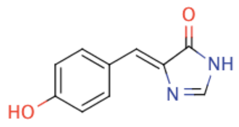 | 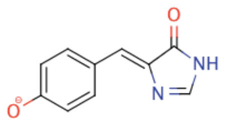 | 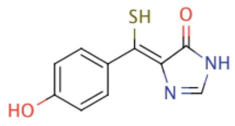 | 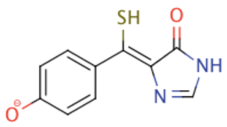 |

**Figure S6:** planar representation of the 8 models for the calculations.

Interestingly, twisting the structure of a non-substituted chromophore has almost no influence (less than 35 nm), while twisting has a major influence on the sulfur-substituted chromophore (170 to 270 nm for anionic and neutral forms, respectively). The absorbance maxima of twisted chromophores are shifted to the red compared to the planar ones, while the intensity of the transition is lowered (smaller *f*). We can conclude in the present study that the covalent bond between the chromophore and the cysteine has the effect of red-shifting the absorbance spectra by both twisting the chromophore and changing the electronic density due to the presence of the sulfur atom.

| Model | Excited state | Transition energy (eV) | Wavelength (nm) | *f* |
|---|---|---|---|---|
| Anionic-$H_{EGFP}$ | S2 | 2.66 | 466 | 0.81 |
| Anionic-$SH_{EGFP}$ | S2 | 2.66 | 467 | 0.64 |
| Neutral-$H_{EGFP}$ | S2 | 3.02 | 411 | 0.60 |
| Neutral-$SH_{EGFP}$ | S1 | 3.00 | 413 | 0.33 |
| Anionic-$H_{AausFP2}$ | S2 | 2.47 | 501 | 0.65 |
| Anionic-$SH_{AausFP2}$ | S1 | 1.93 | 641 | 0.25 |
| Neutral-$H_{AausFP2}$ | S2 | 2.94 | 421 | 0.50 |
| Neutral-$SH_{AausFP2}$ | S1 | 1.80 | 688 | 0.13 |

**Table S1:** Energy of the most intense vertical transition for the different models (in eV), corresponding wavelength (nm) and oscillator strength (*f*).

| | **AausFP1** | **AausFP2** |
|---|---|---|
| **Data reduction** | | |
| Wavelength (Å) | 0.979 | 0.976 |
| Space group | $P2_12_12_1$ | $C222_1$ |
| Cell dimensions | | |
| *a*, *b*, *c* (Å) | 65.08, 101.41, 161.43 | 54.41, 75.13, 100.41 |
| α, β, γ (°) | 90.0, 90.0, 90.0 | 90.0, 90.0, 90.0 |
| Resolution range† (Å) | 50.0 – 2.47 (2.53 – 2.47)* | 50.0 – 2.06 (2.11 – 2.06) |
| Wilson *B*-factor ($Å^2$) | 37.9 | 38.0 |
| No. of reflections | 228,828 (16,791) | 78,841 (5,174) |
| Unique reflections | 39,086 (2845) | 13,079 (976) |
| Multiplicity | 5.9 (5.9) | 6.0 (5.3) |
| Completeness (%) | 99.9 (99.9) | 99.8 (99.7) |
| Mean I/sigma(I) | 9.72 (2.07) | 12.79 (2.16) |
| $R_{meas}$‡ (%) | 21.5 (120.3) | 10.0 (85.1) |
| $CC_{1/2}$ | 0.993 (0.707) | 0.998 (0.683) |
| **Structure refinement** | | |
| Resolution (Å) | 48.42 – 2.47 (2.50 – 2.47) | 44.11 – 2.06 (2.09 – 2.06) |
| $R_{work}$ (%) | 18.1 (29.1) | 16.6 (24.0) |
| $R_{free}$ (%) | 22.8 (35.0) | 20.2 (32.0) |
| No. of atoms | 7567 | 1937 |
| Protein | 7196 | 1788 |
| Solvent | 371 | 149 |
| *B*-factors ($Å^2$) | 37.2 | 24.1 |
| Protein | 37.4 | 22.4 |
| Solvent | 34.1 | 44.3 |
| R.m.s. deviations | | |
| Bond lengths (Å) | 0.010 | 0.009 |
| Bond angles (°) | 1.73 | 1.69 |

†Resolution cutoff based on $CC_{1/2}$.*Values in parentheses are for the outer shell. ‡Rmeas = Rmerge × $[N/(N-1)]^{1/2}$, where N is data multiplicity.

**Table S2.** Data reduction and structure refinement statistics for the structure determination of AausFP1 and AausFP2.

#### Morphological identification

We identified the *Aequorea* individual collected near Heron Island, Queensland using the morphological characters:

- **Bell shape:** Thin and gently convex
- **Diameter of bell:** 30.9mm
- **Number of radial canals:** 49
- **Tentacle pattern:** 3-4 radial canals with no tentacle, then one radial canal with a tentacle
- **Number of tentacles:** 12
- **Gonad shape:** Bilamellar, extending nearly the whole length of the radial canals
- **Stomach size:** Damaged, unknown
- **Tentacle shape:** tentacle bulb tapers adaxially from the margin to a fine point. There is an abaxial keel on each tentacle bulb where the bulb meets the bell margin.

In *Aequorea* the tentacle shape, gonad shape, number of tentacles, and number of radial canals are all strong characters with which to identify a specimen. Overall, the characters listed above closely match the original description of *A. macrodactyla* and further characterizations of the species in later publications. The unique tentacle shape closely matches Kramp's 1961 drawings of *A. macrodactyla* [2]. The number of radial canals the number of tentacles, the bell size, and the tentacle pattern match Kramp's review of *A. macrodactyla* specimens over many size ranges and locales [3]. The shape of the bell and the shape and distribution of the gonads match the original species description [1].

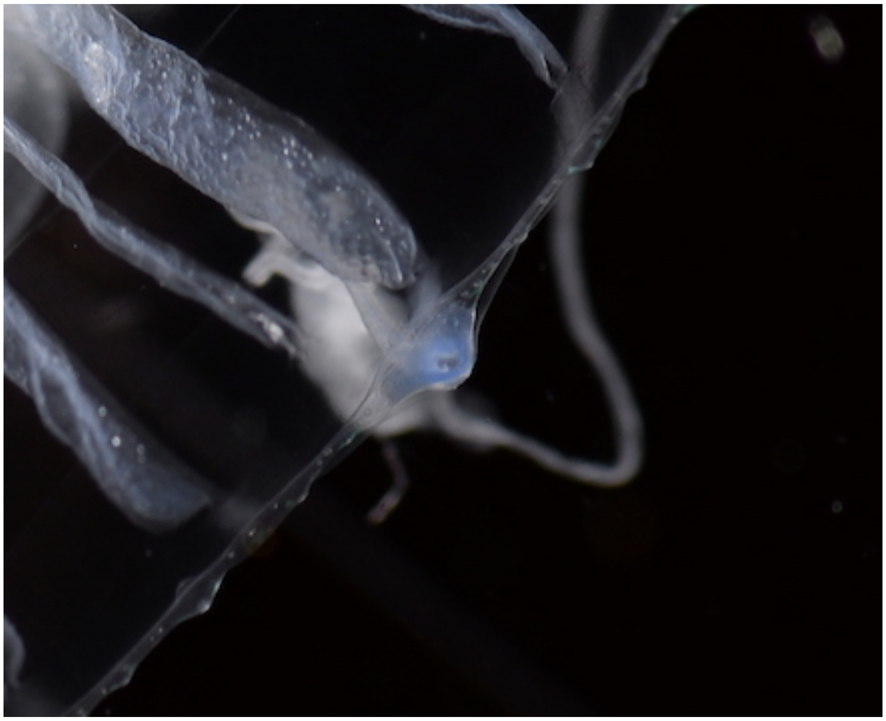

**Fig. S7.** The abaxial keel on each tentacle bulb.

The characters above unambiguously rule out this individual as belonging to sympatric species: *A. australis*, *A. conica*, *A. parva*, *A. globosa,* and *A. pensilis*. The bell of this *Aequorea* specimen is too thin and the gonads are much longer than those found in *A. australis* [15].The bell shape is also too thin, the number of tentacles is too few and the bell width is too large to be either *A. parva* or *A. conica* [16], *A. globosa*, or *A. pensilis*.

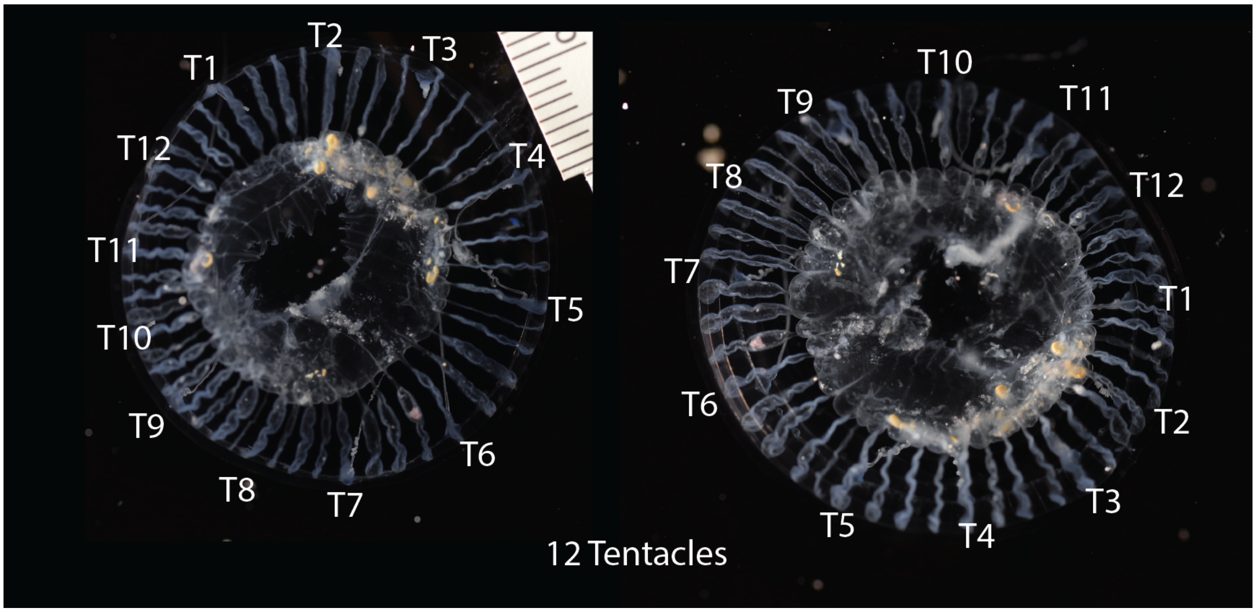

**Fig. S8.** The specimen had 12 tentacles.

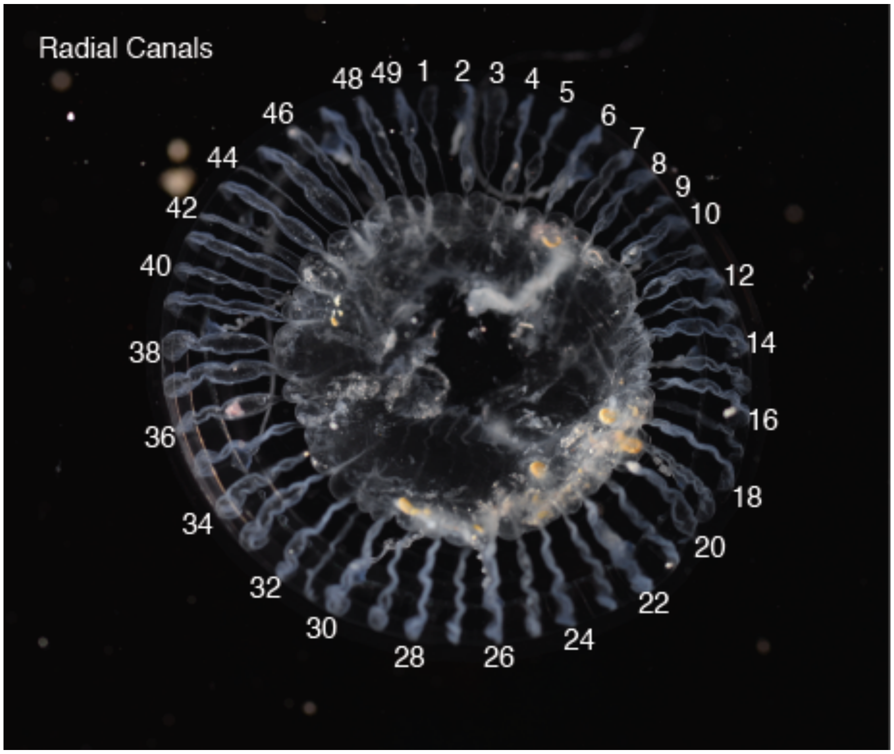

**Fig. S9.** The specimen had 49 radial canals.

#### Molecular identification

We investigated if several DNA barcode sequences, 18S, 16S, and COI, from this *Aequorea* specimen matched the sequences in publicly available databases. We used both a *de novo* assembly and a reference-based assembly to generate the barcode sequences for this specimen.

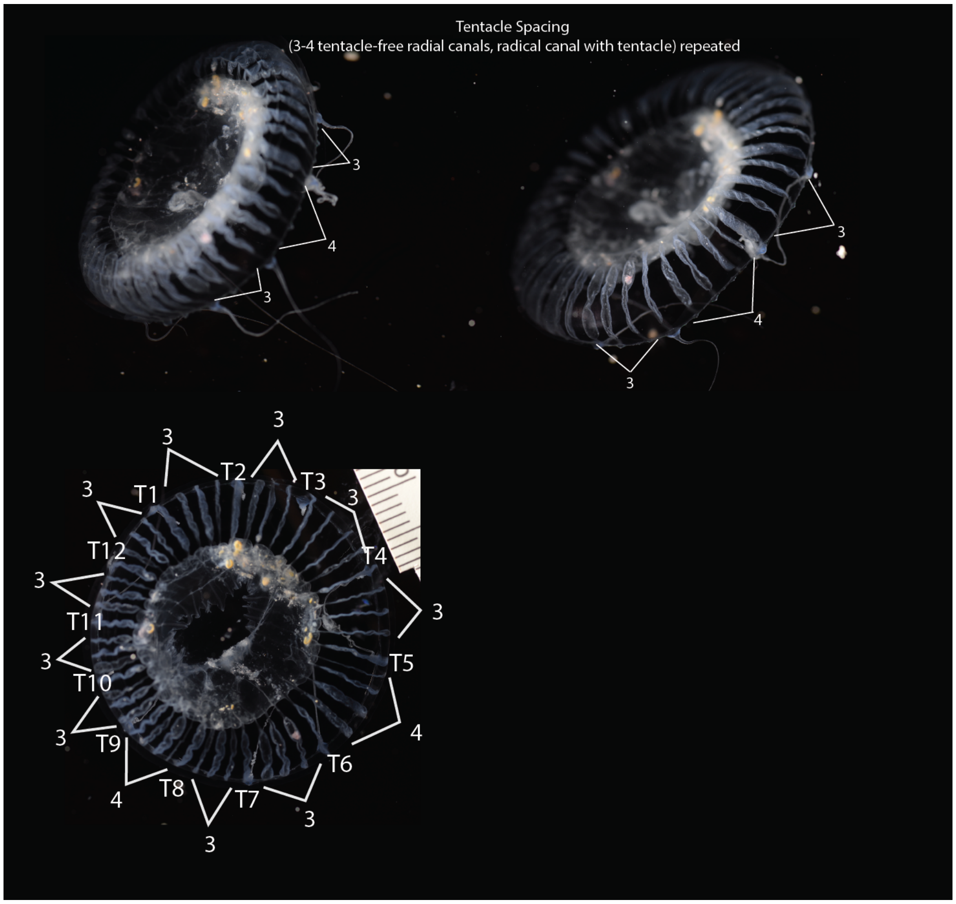

**Fig. S10.** The specimen had 3-4 radial canals between each tentacle.

For the *de novo* assembly approach we used existing *Aequorea* 18S, 16S, and COI sequences as blastn queries against the transcriptome from the Queensland-collected *Aequorea* specimen. We used the following sequences as queries: *A. australis* 18S sequence [KF962202.1](), *A. australis* 16S sequence [KF962390.1](), and *A. macrodactyla* COI sequence [LK054491.1](). The following orthologues were identified in the *de novo* assembled transcriptome:

#### 16S orthologues

```
>TRINITY_DN25108_c0_g1_i1
GTAAATTTTAATTAAAGGTAATTCGTTATAAGTGTGTAAGGCAGGTGGTGAGTTAAAAACCCATTCTAGAGTGTTAGAGGCTTTCAAAGCTTTTACTTAAATATAA
GAAATGGATTTTTGGTGTCTTATTCCTTTCGTACTAAAGACCCTTTAGGTTATTAAACTGTAGATAGAAACCTTCCTGTCTTGCGACGGAATGAACTCAAATCATG
TAAGATTTTAAAGGTCGAACAGACCTACCTTTGATAACTTCTGCATTATCAGGACATCTTAATTCAACATAGAGGTGACAAACTTCATTTTCGATAGGAACTCTAA
AATGAAATTATCCTGTTATCCCTAAGGTAGCTTTTATTTATTGATCGTTTTTCTTTTTACTAAGTAAAAACGGGTCATTATAGCCCATTTTAATGATTGTTAAATG
TATTAATTATACAATAAGTTAAATATTAACGTTGCTTACTTTCGTTAAAATTAGAAAGCAGTCGCCCCAACTAAACTACCAACCTTTTTATTAAGTGAATCAGTTT
TCTTATTATATAAAAAGGTTATAGTTAAGCTCTATAGGGTCTTTTCGTCTTACAGTTTAAAATAGCATCTTAACTATTATTTCAATTTCTATTAATTTTTTCTAAG
ACAGTGGAATATTCGTTCAACCATTCATACTATCCATCAATTAAATGGCGAGTGATTATGCTACATTATCACGGTCAAGGTTACCGCGTCCTTTTAATCTTATTAC
TAAGACTTTACATTATTTAAATTAAAGATCACTGGGCAGGTTACACCTTAGATTTTATCTAGGGCTATGTTTTTGGTAAACAGTCGATATTCTATTTTGCCGAGTT
ACTTAGAAAAAACTCATTACTTTTTTATTCCCTCTTGATTCAGAGTATACAGTCTACAATTCTAATTATTTCTCGAGTATCTTAGATTTATTTCTGTTAGATTGAA
AAGCTGTATAACCTTATTCATGGGTAAAATAAGAAAAATTAGGAACAAATTTAGTAATAATCGTACTTACATTATTTCTTCCGTCAACTTACGAAATCTTTAACGG
AATACTCTACTACTTAAAAATTAGAATTACTGATGAAGAGTTTAACTAAATTTAAGGAGTTAAATCTTTAATTAAATAAACTATTACGTACTCTATAAAATGTGGC
AGTTTCTAAGCCCACTTCTTTGTTGTCATAATAACTAACTCCTTGGCTTTTATCTTCATAAAGTTCATGGATTTCTAATTAAGATTGTTATCTTTTTAACAAATAA
TGTTATAACTATTTGTTTGTACTATTAGAATAATAATTTTACTGATATAAATTTCGAAGAGAACCAGCTATATCCAAGTTCGATTAGTTTTTCGCTGCTAATTGCA
AATCATCTAAGAATTTTTCTACATTCACTAGTTCGTTCATAAAACTGTTCACAATTAGATCACTTGGTTTCGGGAAATATTAACTTAACAGTCTACTGAGCCTAAA
AGCTTGCTAATATTATTTCTTGTAAGTTCATTATACAAAAGGTAAATGATTTTCATGGTTTAACTCTAATCATTGTTAAATTTCCTTCACAGTACTCGTTTTATAG
GTAGATTTAGGTAGTATTTATGTAAAATCATGGTAATTAACATGGAAAAGATTTCGTTCGCCACTACTTTCTTTTTACTCTTATTAGGTTACTAAGATGTTTCAGT
TCTTTTTTAATACTTCAGTTGAAAAATATTAAGTAATTTATTATTATTTAATAATCACTAAACCTCCTATAAATTTGATTAGATTTTTTA
>TRINITY_DN25108_c0_g2_i1
GTAAATTTTAATTAAAGGTAATTCGTTATAAGTGTGTAAGGCAGGTGGTGAGTTAAAAACCCATTCTAGAGTGTTAGAGGCTTTCAAAGCTTTTACTTAAATATAA
GAAATGGATTTTTGGTGTCTTATTCCTTTCGTACTAAAGACCCTTTAGGTTATTAAACTGTAGATAGAAACCTTCCTGTCTTGCGACGGAATGAACTCAAATCATG
TAAGATTTTAAAGGTCGAACAGACCTACCTTTGATAACTTCTGCATTATCAGGACATCTTAATTCAACATAGAGGTGACAAACTTCATTTTCGATAGGAACTCTAA
AATGAAATTATCCTGTTATCCCTAAGGTAGCTTTTATTTATTGATCGTTTTTCTTTTTACTAAGTAAAAACGGGTCATTATAGCCCATTTTAATGATTGTTAAATG
TATTAATTATACAATAAGTTAAATATTAACGTTGCTTACTTTCGTTAAAATTAGAAAGCAGTCGCCCCAACTAAACTACCAACCTTTTTATTAAGTGAATCAGTTT
TCTTATTATATAAAAAGGTTATAGTTAAGCTCTATAGGGTCTTTTCGTCTTACAGTTTAAAATAGCATCTTAACTATTATTTCAATTTCTATTAATTTTTTCTAAG
ACAGTGGAATATTCGTTCAACCATTCATACTATCCATCAATTAAATGGCGAGTGATTATGCTACATTATCACGGTCAAGGTTACCGCGTCCTTTTAATCTTATTAC
```

```
TAAGACTTTACATTATTTAAATTAAAGATCACTGGGCAGGTTACACCTTAGATTTTATCTAGGGCTATGTTTTTGGTAAACAGTCGATATTCTATTTTGCCGAGTT
ACTTAGAAAAAAACTCATTACTTTTTTATTCCCTCTTGATTCAGAGTATACAGTCTACAATTCTAATTATTTCTCGAGTATCTTAGATTTATTTCTGTTAG
```

#### 18S orthologues

```
>TRINITY_DN1762_c0_g1_i1
GAACGAAAGTCGGAGGTTCGAAGACGATCAGATACCGTCGTAGTTCCGACCATAAACGATGCCGACCGGCGATGCGGCGGCGTTATTCCCATGACCCGCCGGGCAG
CTTCCGGGAAACCAAAGTCTTTGGGTTCCGGGGGGAGTATGGTCGCAAGGCTGAAACTTAAAGGAATTGACGGAAGGGCACCACCAGGAGTGGAGCCTGTGGC
>TRINITY_DN24861_c0_g1_i2
ACAAGAAATAACGATACGGGGTCTTTACAGGTCTCGCAATTGGAATGAGTACAATTTAAATCCTTTAACTGTATATAACCGATATTGACTTGACATACATTGACTT
GACTGACATTGACTGTACATGTCTCACGTTGACTGTACATAACTGACTTTCACTGTACATAACTGACGTTGACTGTACATGACTGACATTGACTGTACATGACTGA
TATTTACTTGACTTGGCACAAATTGACTTGACTCTACTAACTTTACATGACTGACGTTGACAGTACATGTCTCACATTGACTGTACATAACTGACTTTCACTGTAC
ATAACTGACGTTGACTGTACATGACTGACATTGACTGTACATGACTGATATT
>TRINITY_DN45236_c1_g1_i1
GATTAAGCCATGCATGTCTAAGTACAAGCCCTTGTACGGTGAAACTGCGAATGGCTCAATAAATCGGTTTTAAGACACTTGATAGTCTCTCTACTTGGATAACCGT
AGCAATTCTAGAGCTAATACATGCTATCCTTCGGGTGAGCTTATTAGATCCAACCACAACCGTGGTGAGTCATAGTAATCAATCTGATTGGCGCAAGCCGACGGTT
CAATCTCGATTCCGCCCTATCAGCTTTCGACGGTAGGGTATTGGCCTACCGTGGCGTTAACGGGTAACGGAGAATTAGGGTTCGATTCCGGAGAGGGAGCCTGAGA
AACGGCTACCACATCCAAGGAAGGCAGCAGGCGCGTAAATTACCCAATCCTAAAGCAGGGAGGTAGTGACAATAAATACTGACGCCGGATTCAATGAATCTGGTAA
AGAATGAGAACAGTCTAAAACCCTTATCGAGGACCCATTGGAGGGCAAGTCTGGTGC
>TRINITY_DN45236_c1_g2_i3
GCGTTGAGGCTATACTACATACTACTACAGTATGTTTTGTACGAATAGTTACCTGGTTGATCCTGCCAGTAGTCATATGCTTGTCTCAAAGATTAAGCCATGCATG
TGTAAGTATAAGCTGTATGTACTGTGAAACTGCGAATGGCTCATTAAATCAGTTATCGTTTATTTGATTGTACTTACCATTAGATGGATATCTGTGGTAATTCTAG
AGCTAATACATGCGAAAAGTCCTGACTCTTCGGGGAAGGGATGTATTTATTAGATTAAAAACCAATGCTCATCTCGGTGGGCTTTAGTGGTGATTCATGATAACTT
TTCGAATCGTATGGGTTTAACCCGACGATGTTTCATTCAAATTTCTGCCCTATCAACTGTCGATGGTAAGGTAGTGGCTTACCATGGTTGTAACGGGTGACGGAGA
ATTAGGGTTCGATTCCGGAGAGGGAGCCTGAGAAACGGCTACCACATCTAAGGAAGGCAGCAGGCGCGGAAATTACCCAATCCCGACTCGGGGAGGTAGTGACAAG
AAATAACGATACGGGGTCTTTACAGGTCTCGCAATTGGAATGAGTACAATTTAAATCCTTTAACGAGGATCCATTGGAGGGCAAGTCTGGTGCCAGCAGCCGCGGT
AATTCCAGCTCCAATAGCGTATATTAAAGTTGTTGCAGTTAAAAAGCTCGTAGTTGGATTTCGGACTGGTGCGTGGTCGGTCTGCCGCAAGGTGTGTTACTGACTC
GTGCTGTCTTCTTCTCAGAGTCCTGCTGTGTACTTAACTGTGCGCAGTATTTTGGAATTTGAGACGTTTACTTTGAAAAAATTAGAGTGTTCAAAGCAGGCTATTT
ATTGCCGGAATACATGAGCATGGAATAATGGAATAGGACTGCGGTCTTATTTTGTTGGTTTCTGAGACCATAGTAATGATTAAGAGGGACAATTGGGGGCATCCGT
ATTTCGTTGTCAGAGGTGAAATTCTTGGATTTACGAAAGACGAACAACTGCGAAAGCATTTGCCAAGAGTGTTTTCATTAATCAAGAACGAAAGTTAGAGGATCGA
AGACGATCAGATACCGTCCTAGTTCTAACCATAAACGATGTCGACTAGGGATCGGCGGGCGTTAAATTTTTAAGATGACTCCGGCGGCACCTTACGGGAAACCAAA
GTCTTTGGATTCCGGGGGAAGTATGGTTGCAAAACTGAAACTTAAAGGAATTGACGGAAGGGCACCACCAGGAGTGGAGCCTGCGGCTTAATTTGACCCAACACGG
GAAAACTCACCAGGT
>TRINITY_DN45236_c1_g2_i4
GATTAAGCCATGCATGTCTAAGTACAAGCCCTTGTACGGTGAAACTGCGAATGGCTCAATAAATCGGTTTTAAGACACTTGATAGTCTCTCTACTTGGATAACCGT
AGCAATTCTAGAGCTAATACATGCTATCCTTCGGGTGAGCTTATTAGATCCAACCACAACCGTGGTGAGTCATAGTAATCAATCTGATTGGCGCAAGCCGACGGTT
CAATCTCGATTCCGCCCTATCAGCTTTCGACGGTAGGGTATTGGCCTACCGTGGCGTTAACGGGTAACGGAGAATTAGGGTTCGATTCCGGAGAGGGAGCCTGAGA
AACGGCTACCACATCCAAGGAAGGCAGCAGGCGCG
>TRINITY_DN45236_c1_g2_i7
GCACCACCAGGAGTGGAGCCTGCTGCTTAATTTGACTCAACACGGGAAAACTCACCAGGTCCAGACATAGTAAGGATTGACAGGTTGAGAGCCCTTTCTTGATTCT
ATGGGTGGTGGTGCATGGCCGTTCTTAGTTGGTGGAGTGATTTGTCTGGTTAATTCCGTTAACGAACGAGACCTTGACCGGCTAAATAGTCGGGCAGTTTTCGAAC
TGTCTCAAT
>TRINITY_DN45236_c1_g3_i1
CCATGGTTGTAACGGGTGACGGATAATTAGGGTTCGATTCCGGAGAGGGAGCCTGAGAAACGGCTACCACATCCAAGGAAGGCAGCAGGCGCGTAAATTACCCAAT
CCTAAAGCAGGGAGGTAGTGACAATAAATACTGACGCCGGATTCAATGAATCTGGTAAAGAATGAGAACAGTCTAAAACCCTTATCGAGGACCCATTGGAGGGCAA
GTCTGGTGC
>TRINITY_DN55608_c0_g1_i1
GCCAGTACTCATATCCTTCTCTCAAACATTAACCCATGCATGTCTAAGAATAAGCTGTATGTACTGTGAAACTGCGAATGGCTCATTAAATCAGTTATAGTTTGTT
TGATGGTACCTGCTACTCGGATAACCGTAGTAATTCTAGAGCTAATACGTGCAACAAACCCCGACTTCTGGAAGGGATGCATTTATTAGATAAAAAGCCGGCGCGG
GCTTGCCCGCGTCTCCGGTGAATCATGGTAACTCGACGGATCGCACGGCCTG
```

#### COI orthologues

No orthologous sequences were identified in the *de novo* assembled transcriptome.

For the reference-based assembly we mapped untrimmed RNA-seq reads to the KF962202.1, KF962390.1, and LK054491.1 sequences using BWA-MEM [17] in unpaired mode. The resulting bam files were used to correct the reference using pilon unpaired mode [18] This same correction strategy was repeated on the output of pilon. The output fasta of the second round of pilon correction was confirmed to be completely converted to the sequence of the Australia-collected *Aequorea* specimen by visual inspection in IGV [19].

#### Phylogenetic analysis

The phylogenetic position of the Australia-collected *Aequorea* specimen was determined using the 16S and COI sequences collected from the transcriptome and the reference-guided assembly using a Bayesian tree. *Aequorea* sp. COI, 18S, and 16S sequences were downloaded from NCBI. ClustalW 2.1 inside Geneious as used to align each locus independently (Cost matrix: IUB, Gap open cost: 10, Gap extend cost: 6.66, Free end gaps) [20,21] Specific sequences can be found in supplemental materials. Alignments were trimmed such that no sequence lacked 5' or 3' sequence. The 18S dataset contained mostly *A. australis* sequences and lacked a significant overlapping region, so we did not proceed with creation of a Bayesian tree.

We produced one Bayesian tree each for COI and 18S using MrBayes [22]. For the 16S tree we used a total chain length of 10000 with four simultaneous chains, a burn-in length of 1000, the HKY85 substitution model with gamma rate variation and a random seed [23]. The COI tree shared the same parameters except it used a chain length of 100000 and a burn-in length of 10000. The COI tree used Nematostella vectensis COI MH087700.1 as the outgroup, and the 16S tree used N. vectensis 16S AY169370.1 as the outgroup.

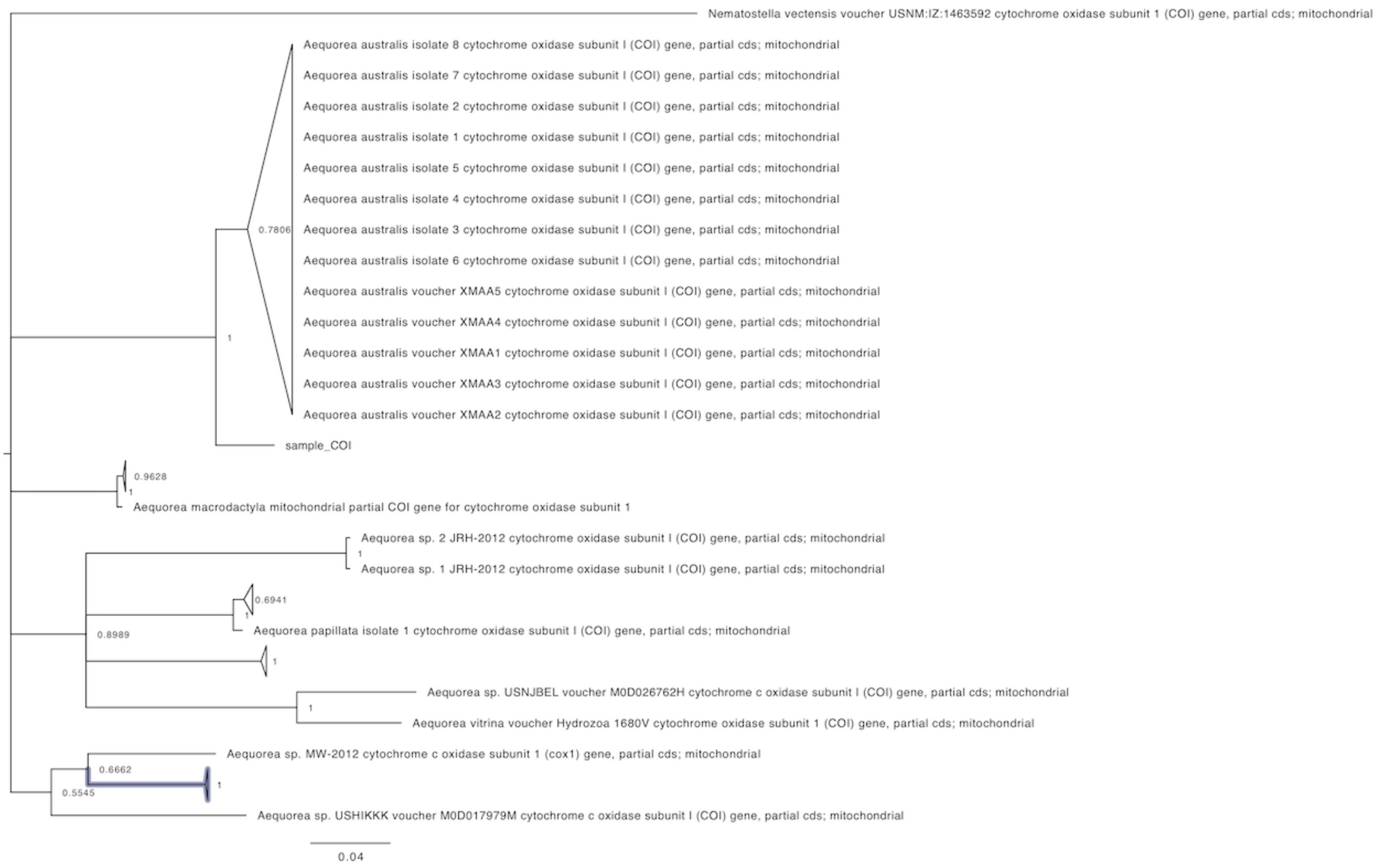

**Fig. S11.** The COI tree shows that the reference-corrected COI sequence (sample_COI) is sister to a large *A. australis* clad. There was no COI sequence identified in the *de novo* transcriptome so there is no comparison. It is important to note that the sample's COI sequence is bona-fide, even though it was assembled using reference-guided assembly. Therefore the sequence coming out as sister to *A. australis* indicates true relation.

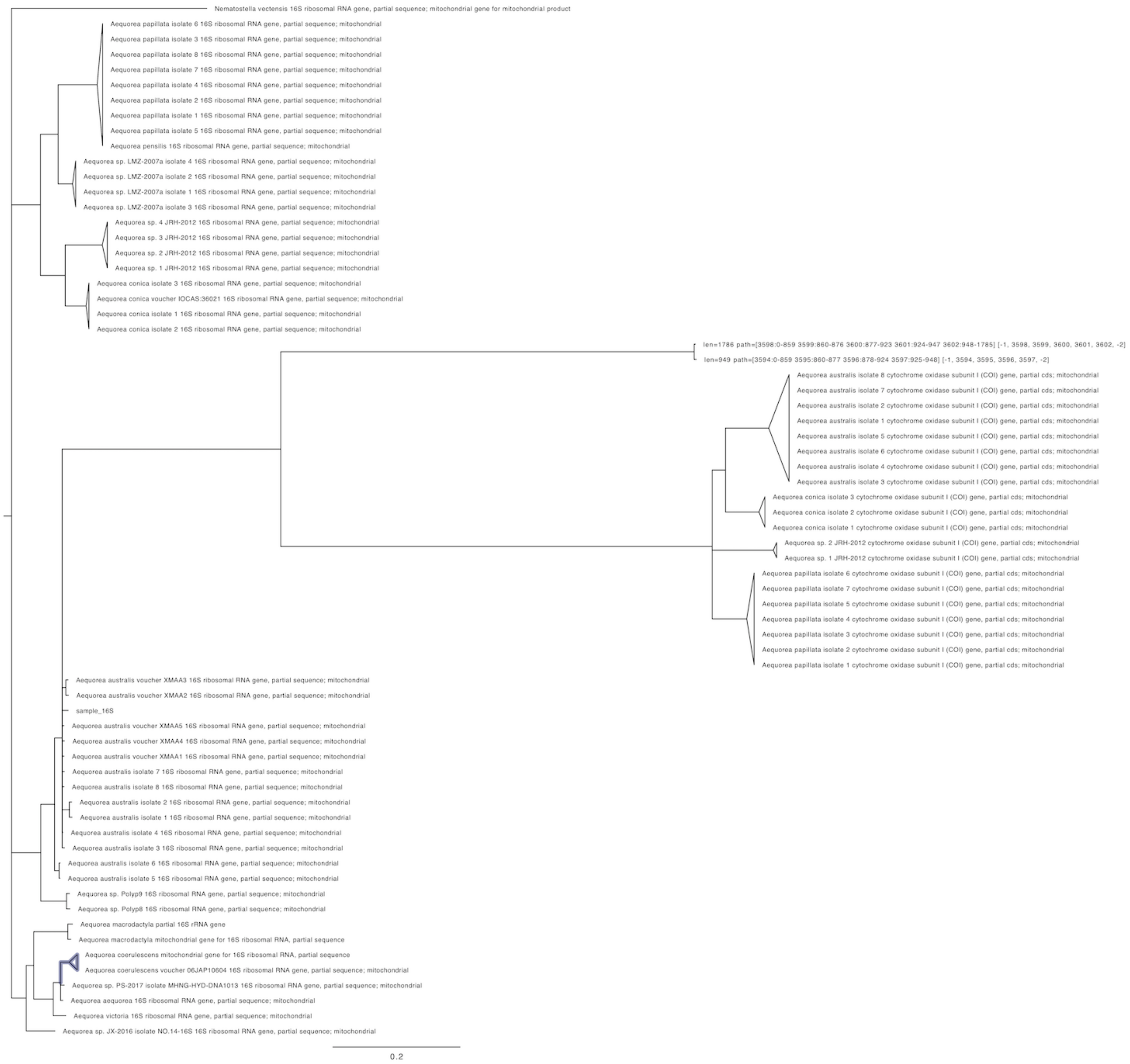

**Fig. S12.** The 16S tree is inconclusive as to the phylogenetic position of both the transcriptomic 16S sequences and the reference-guided assembly 16S sequence. Several species are monophyletic in this tree and *A. australis* is in a large polytomy.

Taken together, the phylogenetic position of the *Aequorea* individual collected near Heron island is unknown, although it shares a similar mitochondrial COI sequence with *A. australis*. As the morphology of the collected individual does not match that of *A. australis* it is possible that there is gene flow between multiple species in the area, causing a *A. macrodactyla* specimen to have an *A. australis* mitochondrial haplotype.

#### AvicFP4 assembly

From the initial *A. victoria* transcriptome assembly we found a transcript that contained a partial GFP open reading frame (ORF).

```
>Av_All_Trinity_TRINITY_DN49895_c4_g3_i2_partial_AvicFP4
ATGGACAGTGGAGCACTTCTTTTCAAGCAAAAAATCCCACTTGTAGTAGAGTTTGAAGGTGATGTCGAGGGAATGAAATTTTCTG
TCAGAGGAAAAGGCCATGGGGATGCAACCAATGGAAGAATTGAAGCCAGCTTTATCTGCACAACTGGAGAATTGCCAGTACCATG
GTCGAGCATTATCAGTAGTCTAACATATGGTTTTCTCTGCTTTACAAAATATCCAGATGGTATCAAAGACTTCCCTAAAAGTGCC
ATGCCAGAAGGATACGTCCAAGAGCGAAAGATTTCTTTTGAAAATGACGGTACATACAAAACACGTGCTGAAGTGACGTTTGAAA
ATGGGTCGGTGTACAATAGAGTCAAGTTGAATGGTGAAGGATTTAACAAAGGTGGAAATATCTTAGGAAAAAAATTGGAATACTC
CTTCAATCCACACTGTATCTACGTTCTTGGAGATAAAGAAAACAATGGCATTAGATGCTGCTTTAATGTTGTTCATAACATTGTT
GGAGGAGGTCAGCTGATTGCCAGCCATAGTCAATTGAATACTCCACTCGGTGGAGGTCCGATAGCTATTCCAGAATACCATCACA
TATGTAACCACACAACACTCAGTAAAGATCCAAGAGAACCACGAGATCACATGACCGTCGC
```

To determine if the partial GFP was a true transcript or not we mapped the RNA-seq reads from the mouth, body (2x libraries), and bell. In total 320 read pairs across the four RNA-seq libraries mapped to this locus. The read depth was drastically inconsistent across the transcript suggesting a misassembly, such as the abrupt increase in coverage around 260 bp, and the lack of 3' coverage (**Fig. S-A**).

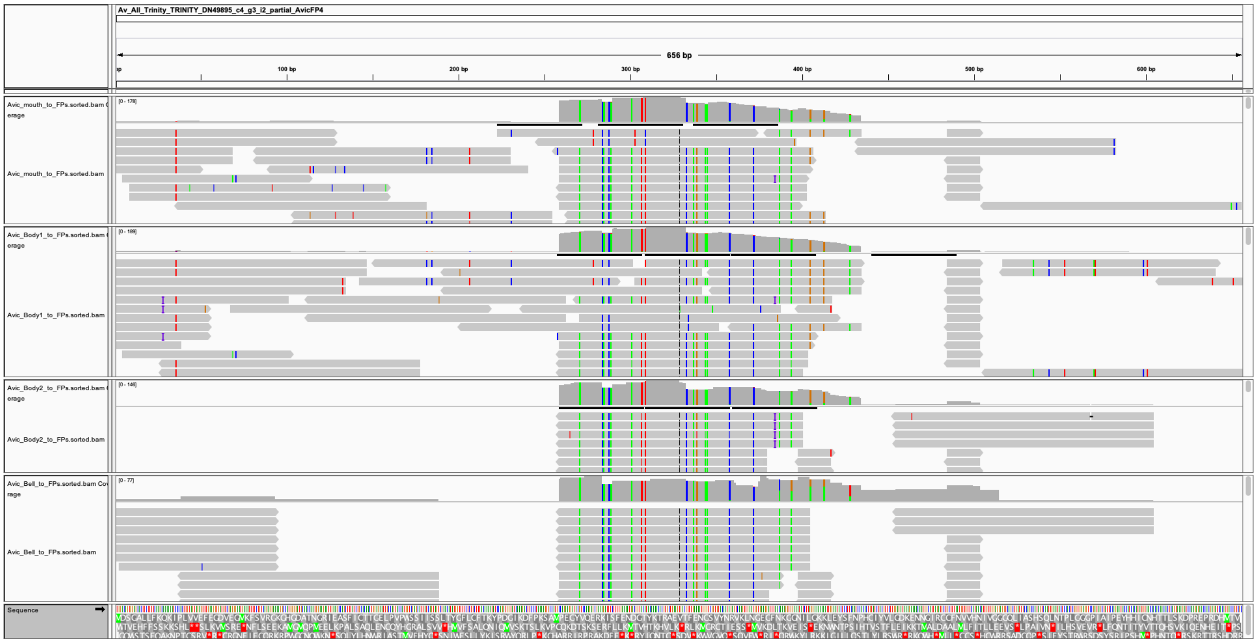

**Fig. S-A.** Reads mapped to the partial *A. victoria* GFP transcript. RNA-seq reads mapped to the partial GFP transcript using BWA-MEM and visualized with IGV. The top track is the mouth RNA-seq library. The middle two tracks are the body RNA-seq libraries, and the bottom track is the bell RNA-seq library.

To attempt to recover a correctly assembled transcript for the partial GFP we used the 320 read pairs mapping to this locus as the input for a *de novo* assembly using the Geneious assembler v11.1.5. One contig contained a complete avGFP homolog ORF. We then mapped all four *A. victoria* RNA-seq libraries to this contig using BWA-MEM to verify it was a real transcript. The transcript was only present in one of the body libraries, and at very low copy number (**Fig. S-B**). After verifying that the protein encoded by this transcript produced a functional FP, we called this transcript AvicFP4. This transcript appears to represent two alleles that differ by one conservative amino acid substitution (Asp to Glu), both of which have nearly identical optical properties. For simplicity, we report here the optical properties of only one of these alleles, and include only one in the sequence alignment in **Fig. 4.**

```
>AvicFP4_complete_transcript
GGAAATCATAAGCGAGACAAGATCATTGTTTTTAATTAATGTTAATTTAAAAGGACCATAATTGAAAAAATTATTAGGGGGCCCC
TAGCCCCCCTGGCTCCGTGGTCCCTGGATTGGTATAAGTACGGCAGAATTTCTATTCGTAAATCCAGCATCAAGAGAACGCACAG
AGCACAGAACATCGAAACCATATATTTACAACAAACTTGAAAAAAAAGATTTTCAAACATGGACAGTGGAGCACTTCTTTTCAAG
CAAAAAATCCCACTTGTAGTAGAGTTTGAAGGTGATGTCGAGGGAATGAAATTTTCTGTCAGAGGAAAAGGCCATGGGGATGCAA
CCAATGGAAGAATTGAAGCCAGCTTTATCTGCACAACTGGAGAATTGCCAGTACCATGGTCGAGCATTATCAGTAGTCTAACATA
TGGTTTTCTYTGCTTTACAAAATATCCAGATGGYATCAAAGACTTCCCTAAAAGTGCCATGCCAGAAGGATACGTCCAAGAKCGA
ACGATCTCTTTTGAAAATGACGGTACATACAAAACACGTGCTGAAGTGACGTTTGAAAATGGGTCGGTGTACAATAGAGTCAAGT
TGAATGGTGAAGGATTTAACAAAGGTGGAAATATCTTAGGAAAAAAATTGGAATACTCCTTCAATCCACACTGTATCTACGTTCT
TGGAGATAAAGAAAACAATGGCATTAGATGCTGCTTTAATGTTGTTCATAACATTGTTGGAGGAGGTCAGCTGATTGCCAGCCAT
AGTCAATTGAATACTCCACTCGGTGGAGGTCCGATAGCTATTCCAGAATACCATCACATATGTAACCACACAACACTCAGTAAAG
ATCCAAGAGAACCACGAGATCACATGACCGTCGCAGAAGTGTTGAAGGCTGTCGATTGCAAGACTGCTTATTTATAGTTCATCAG
ATTTTTTGATGATGTTTTTTCAAGTTTTTGTATTTTAAACATTGCTGCCG
```

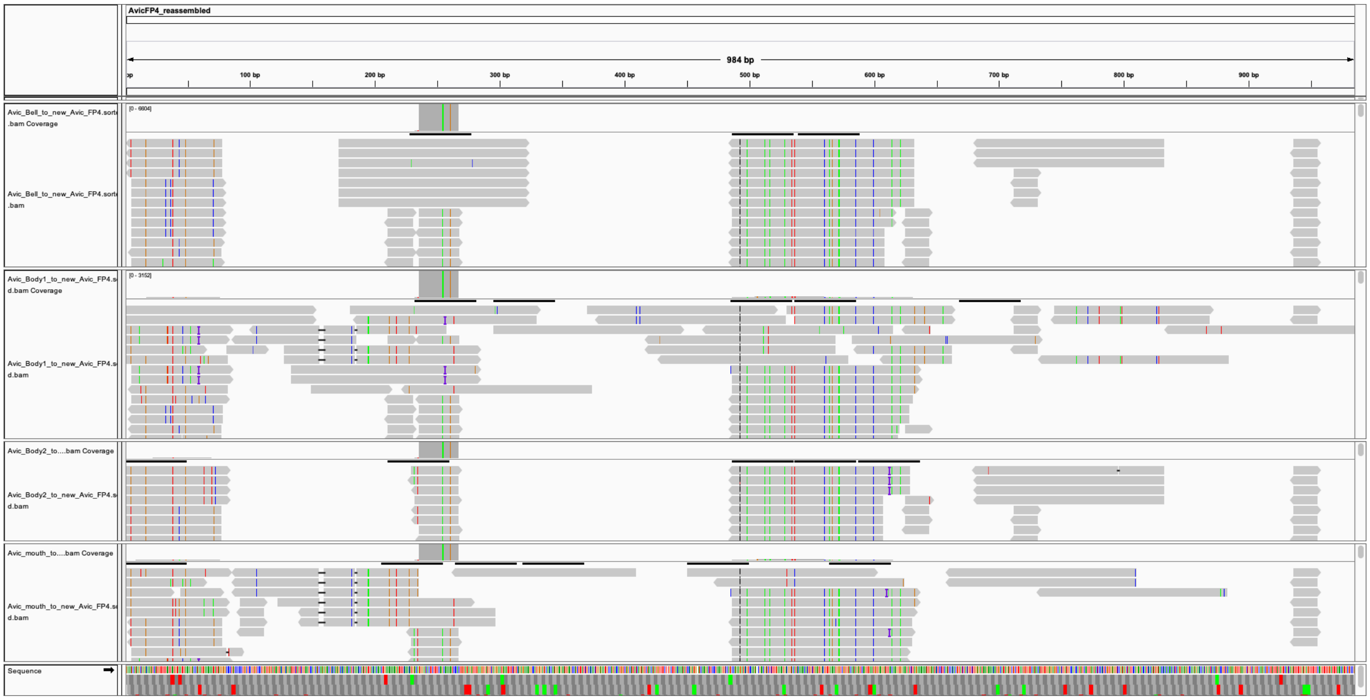

**Fig S-B.** Reads mapped to the *A. victoria* AvicFP4 complete transcript. RNA-seq reads mapped to the partial GFP transcript using BWA-MEM and visualized with IGV. The complete ORF was only expressed in one of the body RNA-seq libraries.

#### Verifying Newly Identified Transcripts

To determine if the remaining newly assembled FP-encoding transcripts from *A. victoria* and *A.* cf. *australis* were *bona fide*, we mapped the RNA-seq reads from each species to their respective transcripts using BWA-MEM. Transcripts AvicFP1, AvicFP2, AvicFP3, AausFP1, AausFP2, AausFP3, and AausFP4 all had continuous read coverage, even in low-copy transcripts. This suggests that all of these transcripts are truly present in the animals (**Fig. S-C** through **Fig. S-I**).

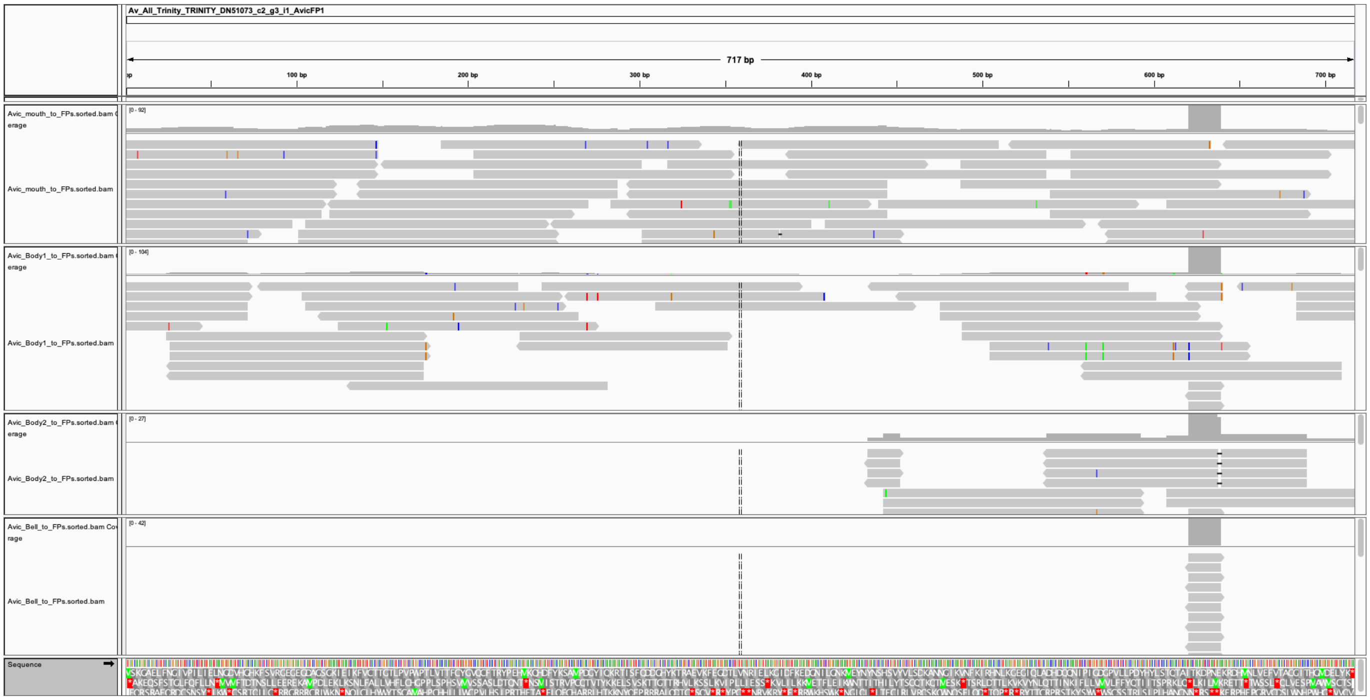

**Fig. S-C:** Reads mapped to the *A. victoria* AvicFP1 complete transcript.

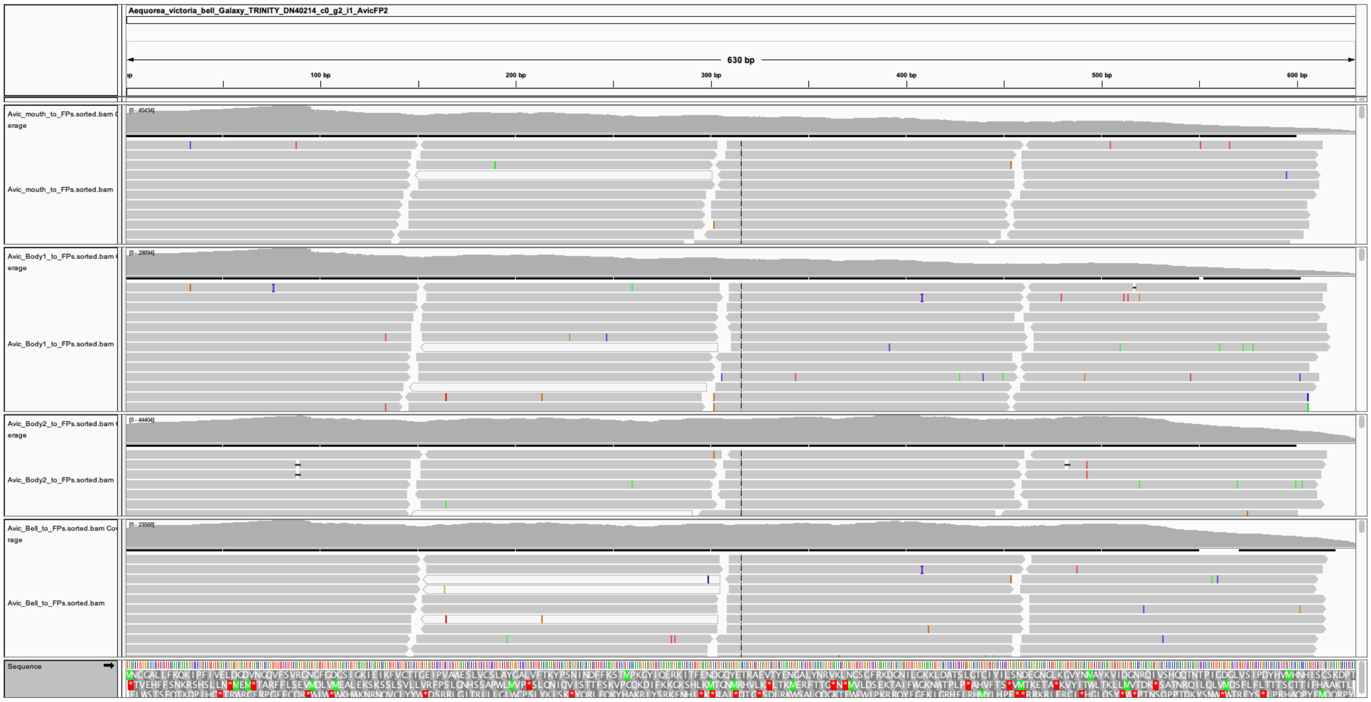

**Fig. S-D.** Reads mapped to the *A. victoria* AvicFP2 complete transcript.

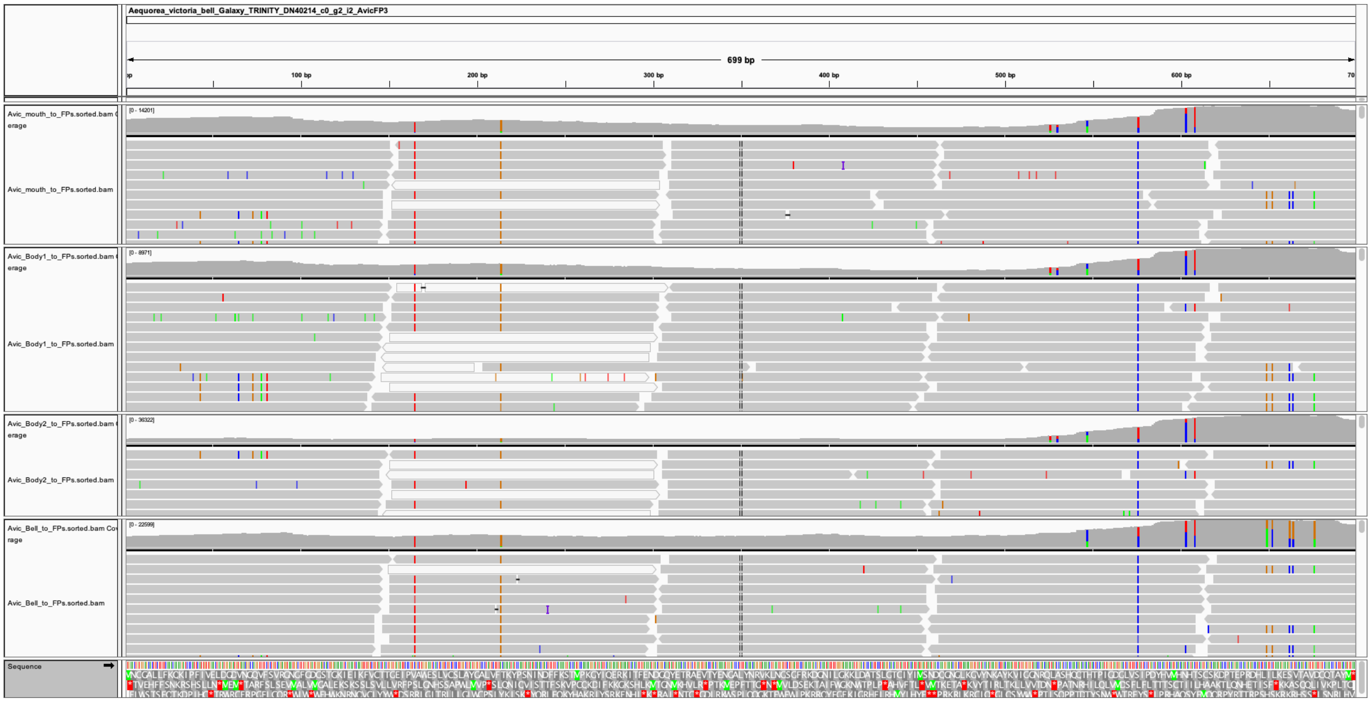

**Fig. S-E.** Reads mapped to the *A. victoria* AvicFP3 complete transcript.

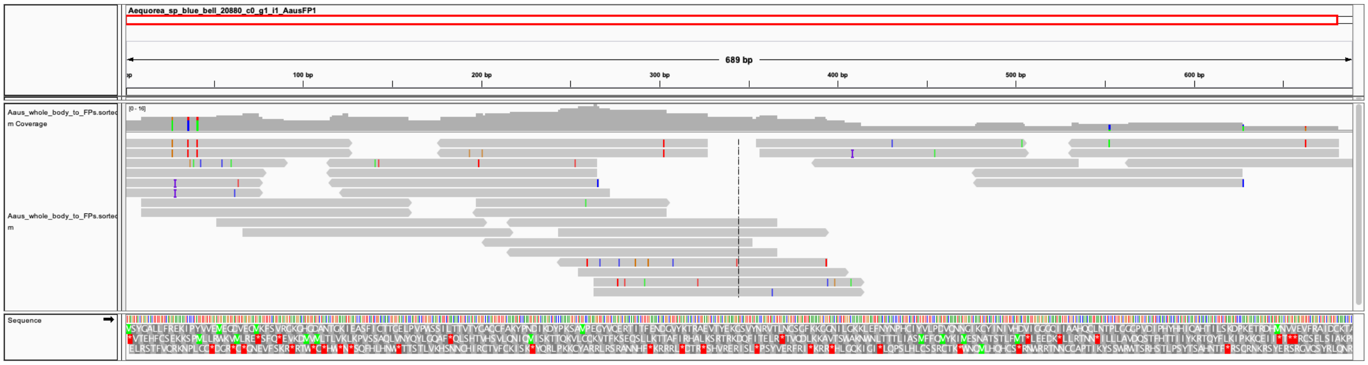

**Fig. S-F.** Reads mapped to the *A.* cf. *australis* AausFP1 complete transcript.

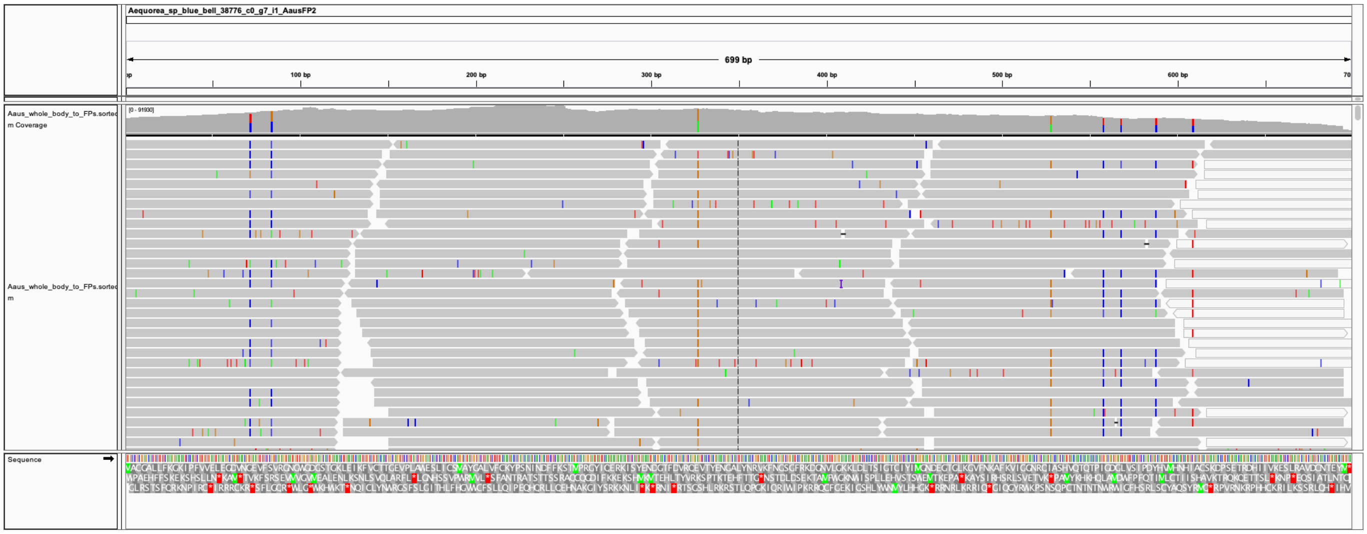

**Fig. S-G.** Reads mapped to the *A.* cf *australis* AausFP2 complete transcript.

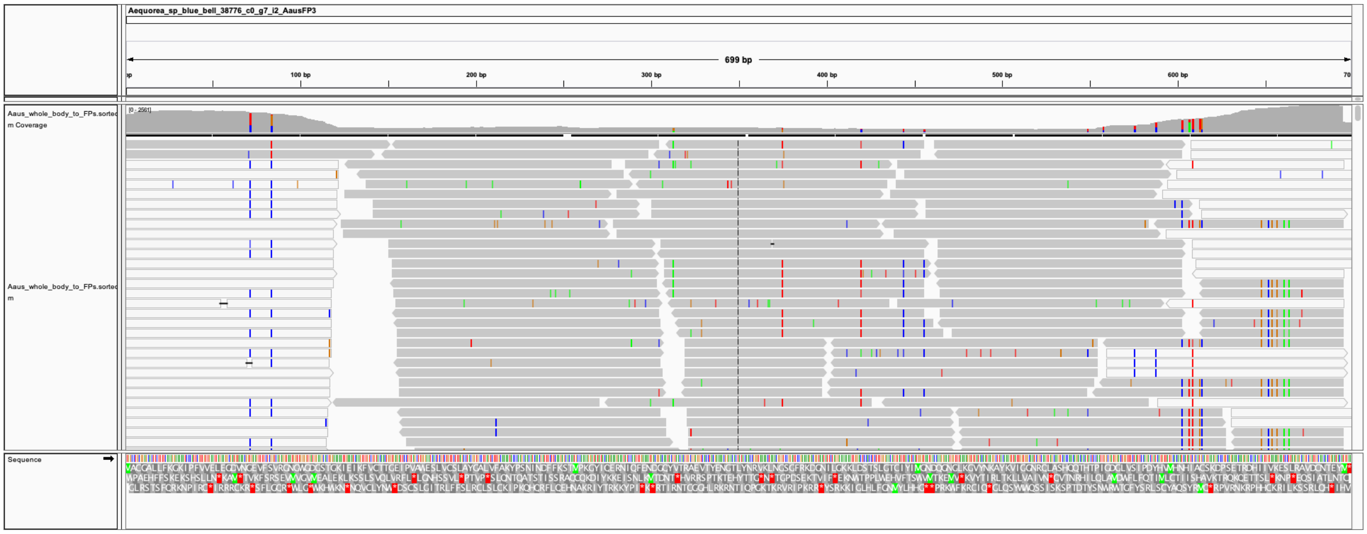

**Fig. S-H.** Reads mapped to the *A.* cf *australis* AausFP3 complete transcript.

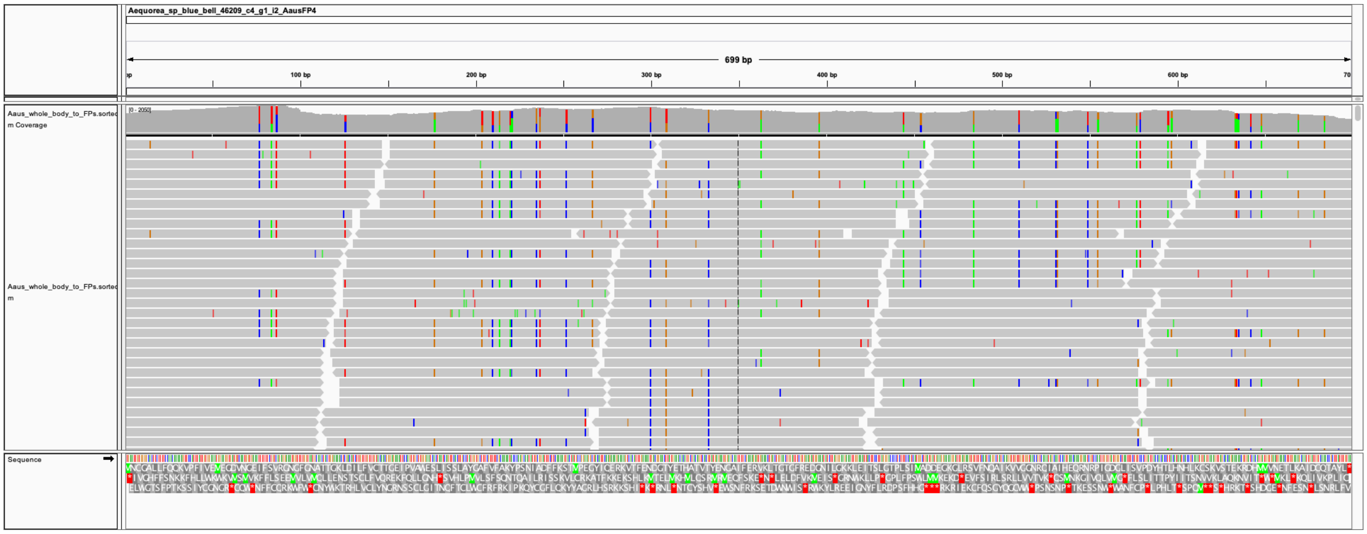

**Fig. S-I.** Reads mapped to the *A.* cf *australis* AausFP4 complete transcript.

### Additional Supplementary Tables and Figures

| | Data on ER Structures | | | | | | | | |
|---|---|---|---|---|---|---|---|---|---|
| | **Condition** | **n** | **mean** | **sd** | **95CI mean** | **median** | **95CI median** | **# normal looking cells (%)** | **# total cells** |
| **1** | mAvicFP1 - D | 166 | **2.18** | 0.85 | **2.05 - 2.31** | 2.03 | 1.88 - 2.24 | 71 | 260 |
| **2** | mAvicFP1 - T | 288 | **2.44** | 0.98 | **2.32 - 2.55** | 2.27 | 2.14 - 2.39 | 61 | 257 |
| **3** | mEGFP - D | 129 | **2.27** | 0.86 | **2.12 - 2.42** | 2.2 | 1.96 - 2.3 | 72 | 212 |
| **4** | mEGFP - T | 27 | **2.07** | 0.7 | **1.78 - 2.35** | 2.07 | 1.6 - 2.38 | 76 | 49 |

**Table S3.** Organized smooth endoplasmic reticulum (OSER) assay results for mEGFP (standard monomer) and mAvicFP1, a novel variant of a previously unknown *A. victoria* green FP. Microscopy was performed by Daphne Bindels for conditions mAvicFP1-D and mEGFP-D, and by Talley Lambert for conditions mAvicFP1-T and mEGFP-T. Numbers in bold represent measured values for the ratio between nuclear envelope and ER intensity.

| AausFP1 | | | |
|---|---|---|---|
| Atom name (chromophore) | Residue name | Atom name (residue) | Distance |
| CD2/CE2 | Leu61 | CD1 | 3.66/3.71 |
| CD1/CG2 | Thr62 | CG2 | 3.79/3.79 |
| CD1 | Ile148 | CD1 | 3.76 |
| CD1 | Ile163 | CD1 | 3.98 |
| CD2/CE2 | Phe222 | CZ | 3.37/3.83 |
| CD2/CG2 | Phe222 | CE1 | 3.47/3.64 |
| OH/CE2 | Tyr143 | CE2 | 3.66/3.95 |
| OH | His146 | CG/CB | 3.45/3.49 |
| CE1 | Ile165 | CD1 | 3.81 |
| OH/CZ | Ala201 | CB | 3.25/3.44 |
| CE2 | Thr203 | OG1 | 3.62 |

**Table S4.** Contacts < 4.0 Å from the AausFP1 chromophore.

| EGFP | | | |
|---|---|---|---|
| Atom name (chromophore) | Residue name | Atom name (residue) | Distance |
| CD1/CG2 | Thr62 | CG2 | 3.69/3.69 |
| OH/CE2 | Tyr145 | CE2 | 3.44/3.92 |
| OH | His148 | CG/CE1 | 3.75/3.81 |
| CE1/CD1 | Val150 | CG2 | 3.75/3.82 |
| CD1 | Phe165 | CE2/CZ | 3.80/3.94 |
| CE1/OH | Ile167 | CD1 | 3.74/3.88 |
| OH/CZ | Thr203 | CB | 3.41/3.66 |
| CZ/CE1 | Thr203 | OG1 | 3.34/3.81 |
| CE1/CZ | Thr203 | CG2 | 3.79/3.85 |
| CE2 | Ser205 | OG | 3.94 |
| CD2/CE2 | Glu222 | CD | 3.84/3.99 |
| CE2/CD2 | Glu222 | OE1 | 3.48/3.51 |
| CD2 | Glu222 | OE2 | 3.80 |

**Table S5.** Contacts < 4.0 Å from the EGFP chromophore.

| AausFP2 | | | |
|---|---|---|---|
| Atom name (chromophore) | Residue name | Atom name (residue) | Distance |
| CE2 | Thr146 | CB | 3.82 |
| OH/CE1 | Asn201 | CB | 3.21/3.31 |
| CE1/CD1 | Asn201 | ND2 | 3.88/3.91 |
| OH | Asn201 | C | 3.20 |
| OH | Asn201 | CA | 3.77 |
| OH | His202 | C/CA | 3.34/3.61 |
| OH/CZ | Ile203 | CG1 | 3.31/3.44 |
| CZ/CE2 | Ile203 | CD1 | 3.19/3.24 |
| CD1/CE1 | Glu220 | CD | 3.43/3.57 |
| CD1/CE1 | Glu220 | OE1 | 3.53/3.88 |
| CD1/CE1 | Glu220 | OE2 | 3.26/3.63 |
| CE1 | Leu222 | CD2 | 4.00 |

**Table S6.** Contacts < 4.0 Å from the AausFP2 chromophore.

| Primer Name | Sequence |
|---|---|
| pNCST-vec-F | CGTTTGATCCGGCTGC |
| pNCST-vec-R | ACCCTGGAAGTACAGGTTTTC |
| AausFP1-F | GAAAACCTGTACTTCCAGGGTATGAGTTACGGAGCACTTTTGTTCAGAG |
| AausFP1-R | GCAGCCGGATCAAACGTCATGCATATGCGGTTTTGCAATC |
| AausFP2-F | GAAAACCTGTACTTCCAGGGTATGGCCTGCGGAGCACTTCTTTTC |
| AausFP2-R | GCAGCCGGATCAAACGTCACATGTATTCAGTGTTGCAATCGACTG |
| AausFP4-F | GAAAACCTGTACTTCCAGGGTATGAATTGTGGGGCACTTCTTTTC |
| AausFP4-R | GCAGCCGGATCAAACGTCACAAATAAGCGGTTTGACAATCAATTG |
| AvicFP1-A206K-F | CCCAGACCAAGATCACCAAGGACCCTAACG |
| AvicFP1-A206K-R | GGTCCTTGGTGATCTTGGTCTGGGTAGACAGG |

**Table S7.** Primers used in this study.

| Species | Protein | Bell Margin | Body | Mouth |
|---|---|---|---|---|
| *A. victoria* | avGFP | 38 | 1.3 | 0.7 |
| | AvicFP1 | 0.0 | 0.3 | 0.0 |
| | AvicFP2 | 0.0 | 1.7 | 0.0 |
| | AvicFP3 | 0.0 | 0.4 | 0.0 |
| | AvicFP4 | 8.4 | 3.7 | 6.6 |
| | | **Whole Animal** | | |
| *A.* cf. *australis* | AausFP1 | 0.1 | | |
| | AausFP2 | 520 | | |
| | AausFP3 | 4.6 | | |
| | AausFP4 | 22 | | |
| | AausGFP | 2.3 | | |

**Table S8.** Transcript abundance for FPs in this study, shown as a percent abundance relative to the housekeeping gene GAPDH. Because we sampled only one animal of each species, it is impossible to estimate the error in these data or variance between individuals since only one animal was sampled for each species, but it is clear that many FP transcripts were present at very low levels in the sampled animals—notable examples include all chromoproteins from *A. victoria* and the extremely bright AausFP1 from *A.* cf. *australis,* which is over 5000-fold less abundant than the chromoprotein AausFP2.

Chain A

Chain B

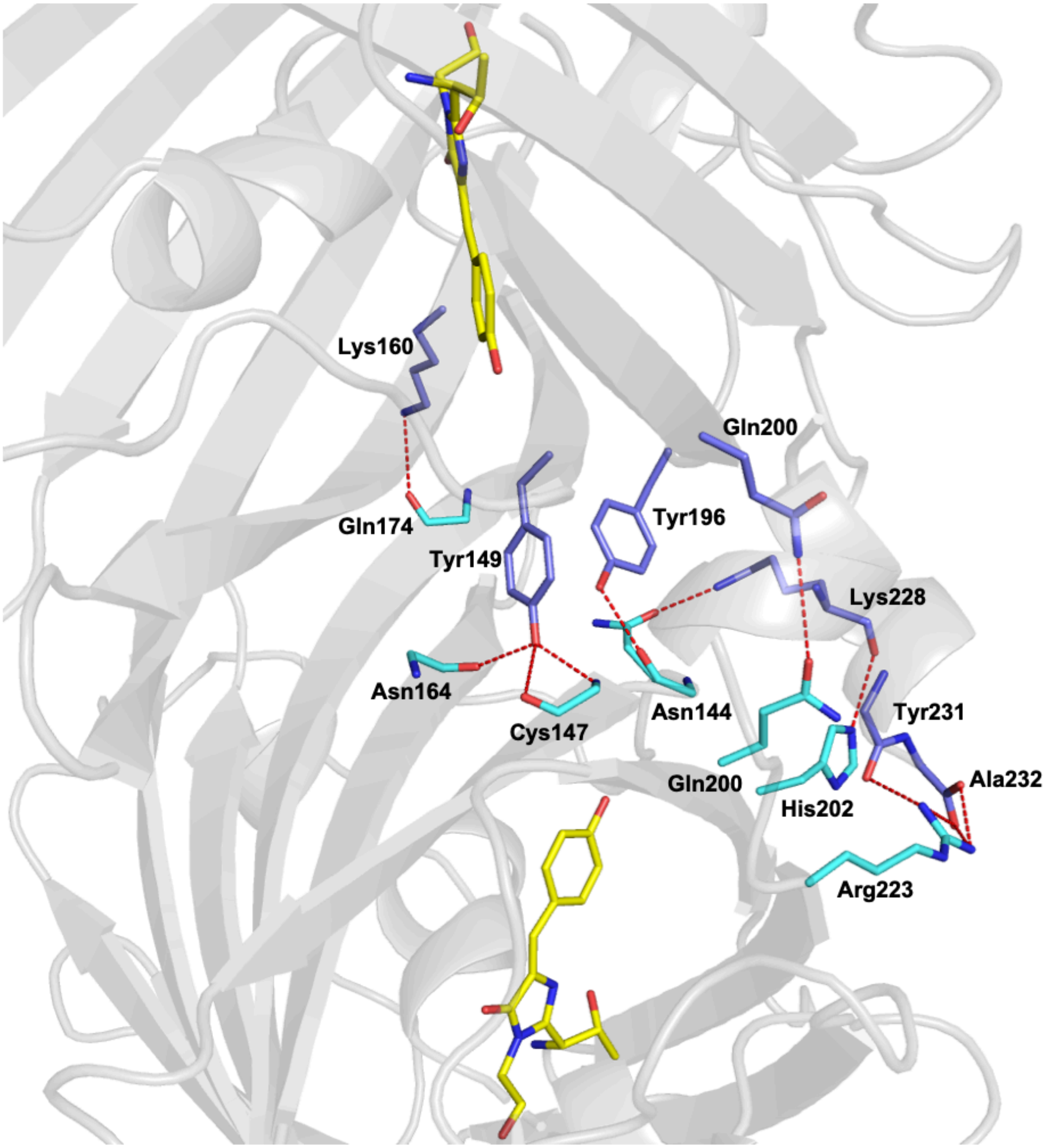

**Figure S13.** A diagram of the critical interactions comprising the AausFP1 dimer interface.

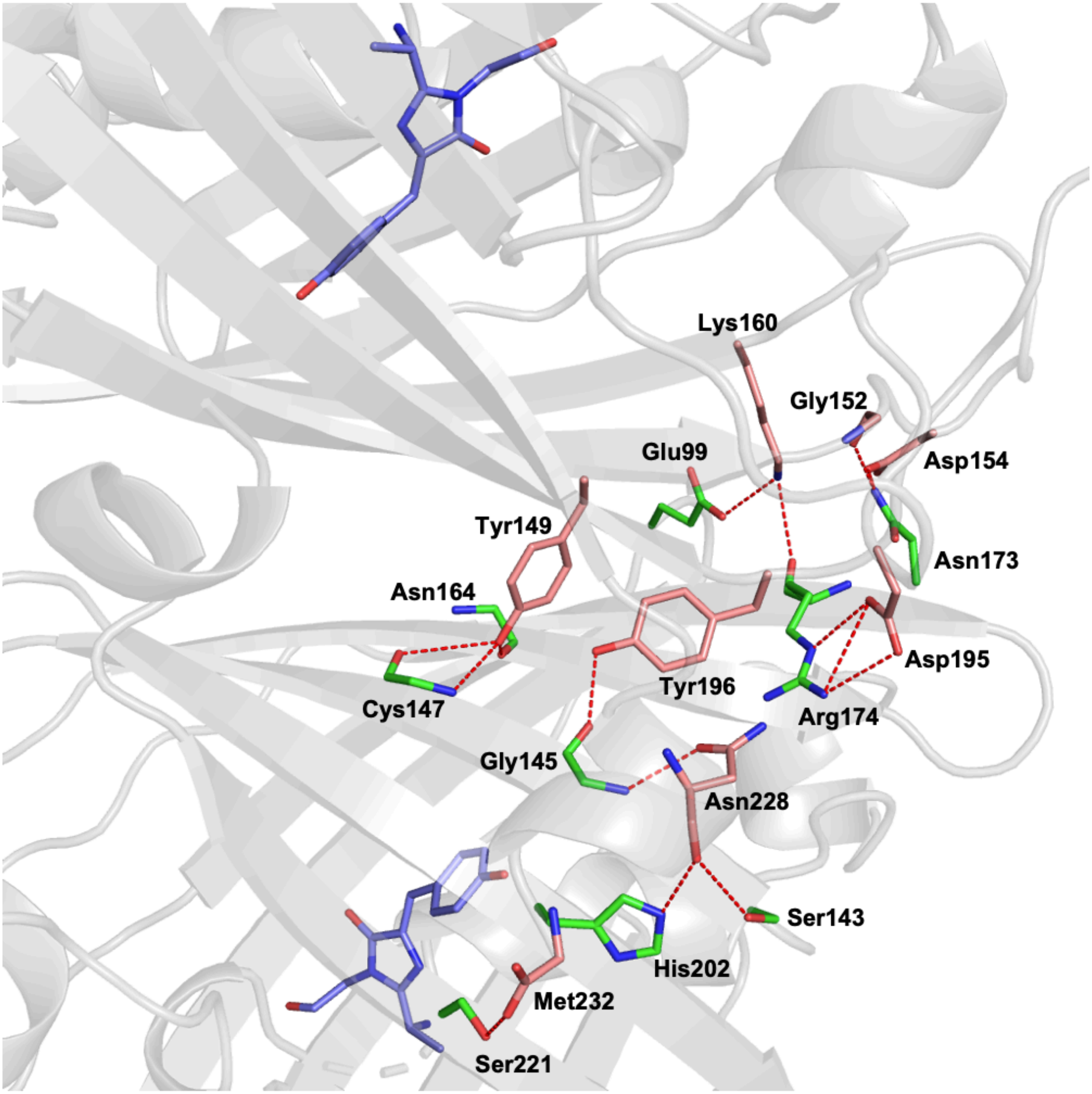

**Figure S14.** A diagram of the critical interactions comprising the AausFP2 dimer interface.
